# Ancient species diversity and niche adaptation in *Tannerella* and *Porphyromonas* revealed through pangenomics

**DOI:** 10.64898/2026.02.09.704811

**Authors:** Aurore Galtier, Christina Warinner, Irina M. Velsko

## Abstract

*De novo* assembly of ancient and modern bacterial metagenomes can shed light on evolution and ecology of bacterial species that are challenging to culture. *Tannerella* and *Porphyromonas* are bacterial genera linked to periodontal disease, and understanding their evolution may reveal insights into their role in oral disease development. We performed pangenomic and phylogenetic analyses on a global set of isolates and metagenome-assembled genomes of the genera *Tannerella* (n=238) and *Porphyromonas* (n=976), including 66 genomes from ancient dental calculus samples (up to 14,800 years old), and modern oral samples from present-day living populations. We identify a novel species of oral *Tannerella* in modern and ancient humans, which we call Ca. *Tannerella abscondita*, that is related to and often mistaken for *Tannerella forsythia* but differs in its virulence repertoire. We reveal distinct niche tropism in *Tannerella* species and *Porphyromonas pasteri,* but not *Porphyromonas gingivalis*. There is limited phylogeographic structuring, and virulence genes are homogeneously distributed across continents and oral niches. Saliva-derived strains of *T. forsythia* and *P. gingivalis* from Oceania and *T. serpentiformis* and *P. pasteri* from Asia show enrichment of pseudogenes related to ecological niche transitions. A phylogenetic analysis of the *P. gingivalis* major fimbrial protein gene *fimA* reveals the genes cluster by genotypes, and that no ancient genes are found in genotypes I and Ib. Using *de novo* assembly for bacterial pangenomics improves the representation of oral genera found in reference databases and enhances our ability to study the evolutionary history of these taxa.

**Significance statement:** Pangenome analyses have been primarily applied to clinically significant species that are abundant and easy to culture, leaving a substantial number of human commensal microbes uninvestigated. *De novo* assembly of commensal microbial metagenomes is a significant resource for generating metagenome assembled genomes (MAGs) of difficult-to-culture species, which can be used for pangenome analyses. We investigated the ecological niches and geographic localization of multiple species of the human oral commensal genera *Tannerella* and *Porphyromonas* using MAGs assembled from globally diverse modern and ancient populations. While much work on these genera comes from Europe and North America, we note distinctions in ecological niches and species prevalence globally, highlighting how MAGs overcome limitations set by culture-based approaches that dominate currently available data. Additionally we demonstrate how improved taxonomic representation of diverse oral species through MAGs can clarify the evolutionary history of commensals and their virulence factors associated with oral disease.

## Introduction

Co-evolution of resident gut bacteria and the host has been shown in humans and other primates(Sanders et al. 2023; Suzuki et al. 2022) indicating co-diversification of commensal microbes with their host. Whether the commensal bacteria in the mouth show similar evidence of co-diversification and sensitivity to host behavior has been investigated to only a limited extent(Velsko et al. 2026; Fellows Yates et al. 2021), and appears to show a similar trend. Oral microbial communities are more stable than gut communities to sudden short-term perturbations such as antibiotic exposure(Zaura et al. 2015), and have been maintained by primate hosts for millennia(Velsko et al. 2026; Fellows Yates et al. 2021), making them intriguing candidates for the study of co-evolution of commensal species with the human host(Fellows Yates et al. 2021).

Little is known about the co-evolution of most oral bacterial species with the host because research tends to focus on a small number of medically relevant taxa(Jensen 2025). Now, with the increasing feasibility reconstructing oral genomes from ancient dental calculus (calcified dental plaque biofilms preserved on tooth surfaces in archaeological skeletons), deep-time investigation of oral microbial evolution is possible using historic and ancient oral microbiome samples up to 102,000 years old(Fellows Yates et al. 2021). We recently showed that genes involved in carbohydrate processing distinguish oral *Streptococcus* species in humans from other primates, suggesting commensal microbial genomic adaptation to human dietary developments through human evolution(Fellows Yates et al. 2021). Evidence of additional dietary carbohydrate adaptation comes from *Streptococcus mutan*s, a species strongly associated with dental caries, which appears to have expanded its repertoire of carbohydrate processing genes concomitant with the advent of agriculture in Europe(Cornejo et al. 2013), supporting that oral microbes are sensitive to host behavior.

Advances in the *de novo* assembly of globally and temporally diverse oral sample metagenomes are now substantially expanding the known diversity of oral microbes from ancient and living populations across a range of lifestyles(Velsko et al. 2026). Similar to what is seen with gut microbiota, a number of taxa are negatively correlated with high levels of industrialization and urbanization, with a large diversity in ancient calculus that is missing in modern industrial populations but still found in non-industrial populations(Velsko & Warinner 2025; Velsko et al. 2026). We determined that current reference databases such as NCBI lack representation of taxonomic and genomic diversity found in oral taxa, and therefore incorporating metagenome-assembled genomes (MAGs) from diverse sources, both modern and ancient, into genomics studies should provide a more globally comprehensive view of oral microbial population ecology and evolution.

Of particular interest in oral health research are taxa involved in periodontal disease, which results from microbiota-mediated inflammation of soft tissues and bone supporting the teeth (Baker et al. 2024; Lamont & Hajishengallis 2015). Periodontal disease has historically been associated with three oral bacterial species, collectively termed the “red complex”(Socransky et al. 1998): *Porphyromonas gingivalis*, *Treponema denticola*, and *Tannerella forsythia*. All three are anaerobic and colonize dental biofilms late in biofilm development. No large-scale, global pangenomic investigations of these species exist, in part due to few available genomes in databases such as NCBI’s Genbank(Suresh et al. 2025; Zwickl et al. 2020; Stocke et al. 2024; Miranda-López et al. 2024). Therefore, questions regarding the extent of genetic and genomic diversity present in these species today, whether the pathogenic potential of strains is evenly globally distributed(Hilbig et al. 2026), and how gene content has changed over time, remain unanswered. We recently conducted a global biodiversity study of >20,000 oral microbial genomes obtained from the published literature and from newly assembled metagenomes. The abundance of *Porphyromonas* and *Tannerella* MAGs in this dataset(Velsko et al. 2026) provided the opportunity to explore these outstanding questions.

Here we explore and refine the diversity of the genera *Tannerella* and *Porphyromonas* in oral samples spanning multiple continents, time periods, and lifestyles, as related to reference genomes. We identify phylogeographic adaptation and oral biogeographic partitioning in specific species, and report three new *Tannerella* species in humans, for which we propose the names Candidatus *Tannerella abscondita*, Ca. *T. ophioides*., and Ca. *T. anguiformis*. We find that *T. abscondita* was previously described but mistakenly identified as a lineage of *T. forsythia* in ancient dental calculus metagenomes. Combining MAGs with reference genomes enables new insights when studying the evolutionary history of oral microbiota.

## Results

### MAGs reveal extensive unexplored diversity in Tannerella and Porphyromonas

From a collection of 24,878 oral MAGs of medium and high quality(Velsko et al. 2026), we identified 210 that were classified as *Tannerella* and 781 that were classified as *Porphyromonas*, including 48 *Tannerella* and 18 *Porphyromonas* MAGs derived from ancient dental calculus. To clarify the number of species represented by the MAGs, their relation to published genomes in these genera, and where ancient MAG diversity fits into present-day diversity, we downloaded all available *Tannerella* (n=28) and *Porphyromonas* (n=195) genomes in NCBI that were classified in these genera by the Genome Taxonomy Database (GTDB) r214 to curate a large and comprehensive collection of genomes (Supplemental Figures S1, S2, Supplemental Tables S1, S2). The NCBI isolate genomes and MAGs come from human samples from Europe, Africa, Asia, Oceania, South America, and North America, representing broad global and lifestyle diversity, as well as from present-day dog oral samples. The dental calculus MAGs span 15,000 years of human history, and also include MAGs from chimpanzees, gorillas, howler monkeys, and brown bears (Figure 1). The ancient MAGs are of comparable quality to MAGs from modern-day oral samples (Supplemental Figures S3-S5, S8, S9, Supplemental Table S1, S2).

**Figure 1:**
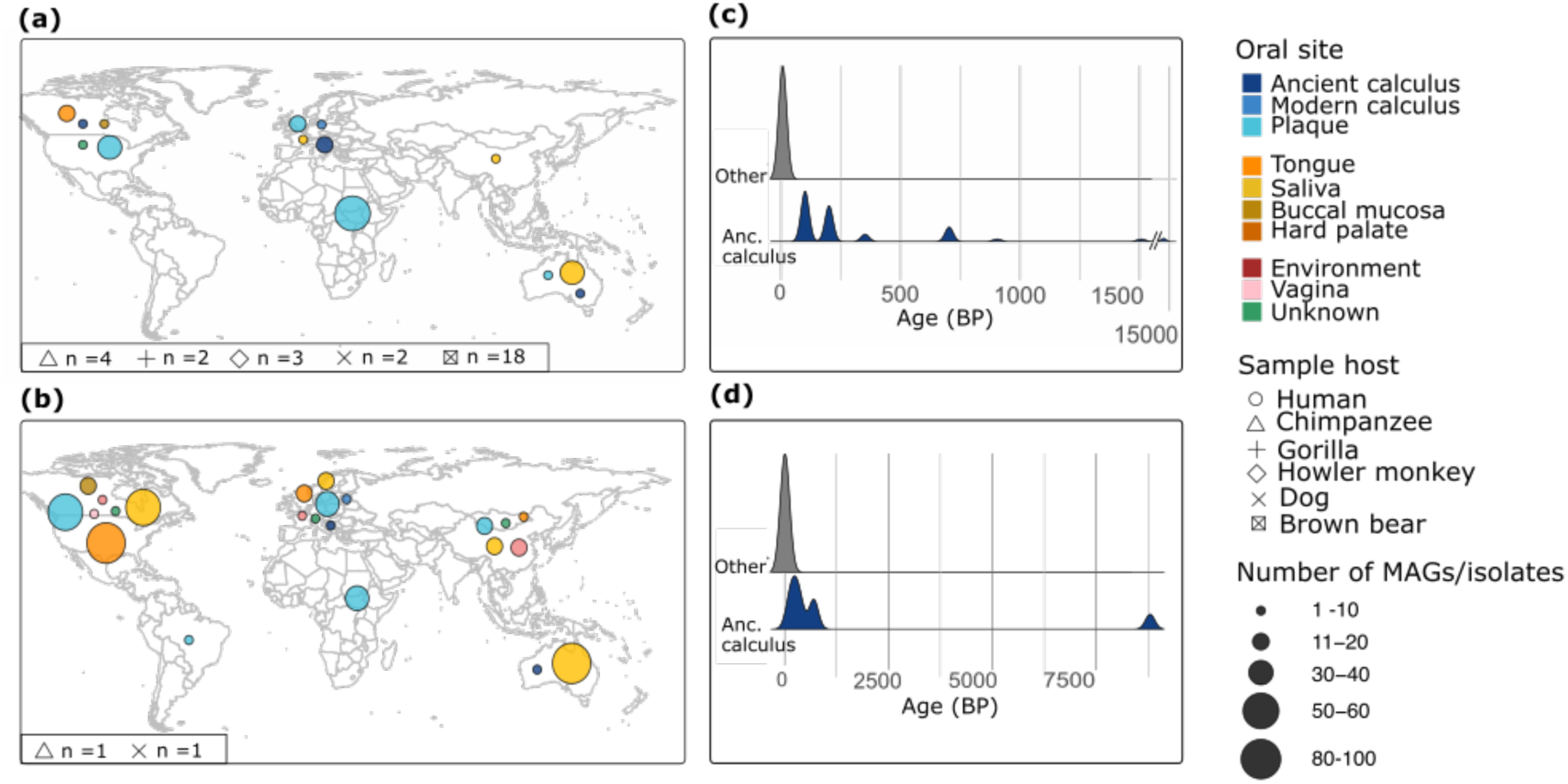
Geographic origin of MAGs and reference genomes of *Tannerella* and *Porphyromonas.* **(a)** Map of the distribution of human-derived *Tannerella* genomes and MAGs by continent. Counts of non-human-derived MAGs are at the bottom of the panel. **(b)** Map of the distribution of human-derived *Porphyromonas* genomes and MAGs by continent. Counts of non-human-derived MAGs are at the bottom of the panel.All chimpanzee, gorilla, howler monkey, and brown bear MAGs come from ancient dental calculus off museum specimens ca. 200 years old; all dog MAGs are from present-day samples. **(c, d)** Histogram showing the time range of human-derived MAGs in years before present (BP). Anc. calculus - ancient calculus; Other - all modern oral sample types. **(c)** Time scale of *Tannerella* genomes and MAGs. **(d)** Time scale of *Porphyromonas* genomes and MAGs. Color corresponds to the host oral source.

To determine the number of species in our datasets, we clustered all genomes and MAGs by genus (238 *Tannerella* genomes; 976 *Porphyromonas* genomes) based on their average nucleotide identity (ANI). Using a cut-off of 95% ANI to define species clusters, we identified 14 species clusters in *Tannerella* and 61 species clusters in *Porphyromonas*. Hierarchical clustering of genomes based on ANI indicated the *Tannerella* species clusters fell into two groups at an ANI of 65% (Supplemental Figure S1), with ancient genomes found in 10 species clusters across both higher-order groups. We chose to assess the diversity of all *Tannerella* species clusters based on these 65% ANI groups, here termed the *T. forsythia-*group (six species clusters) and the *T. serpentiformis*-group (eight species clusters), based on the only named species cluster found in each group. Comparing clustering of the *Porphyromonas* genomes based on ANI indicated higher genomic diversity within this genus (down to 50% ANI, Supplemental Figure S2), and we chose to focus on the two species clusters with >100 genomes, both of which include ancient MAGs: one cluster representing *P. gingivalis* (189 MAGs/genomes) and the other representing *P. pasteri* (315 MAGs/genomes), two recognized oral species. We focused on the ANI-defined species clusters selected above to assess the phylogenetic relationships within and between species clusters, their pangenome content, and their virulence genes.

### Phylogenetic relationships of oral Tannerella

First we investigated genomic and genetic diversity in the genus *Tannerella*, which includes only three species with formal names in the NCBI database: *Tannerella serpentiformis, Tannerella forsythia*, and *Tannerella pachnodae*. Of these, *T. forsythia* and *T. serpentiformis* are oral commensals of humans, while *T. pachnodae* is a gut commensal of insects. Our oral *Tannerella* MAGs represent *T. forsythia*, *T. serpentiformis*, and 12 unnamed species clusters based on ANI clustering. To understand the phylogenetic relationship between these species clusters, we built two independent marker gene-based maximum-likelihood phylogenies, one for each of the two 65% ANI clusters (Figure 2a,c). We found that the 95% ANI-determined species clusters correspond to individual phylogenetic clades (Figure 2b,d, Supplementary Figure S3), supporting their designation as new species clusters. These phylogenies recapitulate the MAG- and reference genome-based *T. forsythia* tree that we built for an independent study(Velsko et al. 2025), and confirm our previous interpretation that additional species beyond *T. forsythia* are present in humans, as well as other host species.

**Figure 2:**
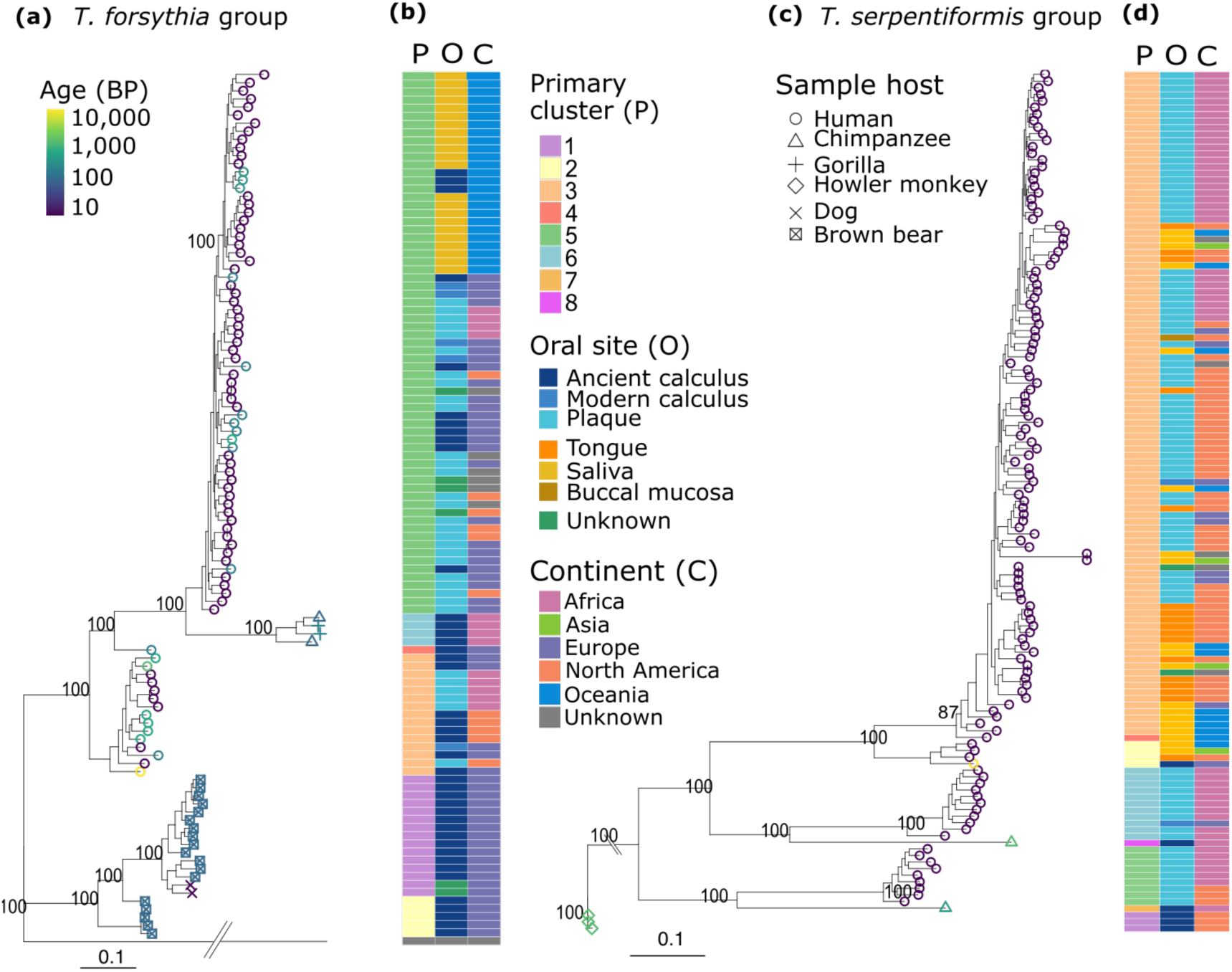
Phylogenetic relationship of *Tannerella* species clusters to geographic origin and oral sample site. **(a)** Maximum-likelihood phylogenetic tree of the *Tannerella forsythia*-group; *T. serpentiformis* was used as the outgroup to root the tree. **(b)** Metadata matrix of the *Tannerella forsythia*-group. **(c)** Maximum-likelihood phylogenetic tree of the *Tannerella serpentiformis*-group; a *T. serpentiformis*-group genome from a howler monkey was used as the outgroup to root the tree. **(d)** Metadata matrix of the *Tannerella serpentiformis*-group. Tree tip color corresponds to sample age in years before present (BP) and tip shape corresponds to sample host. The matrix gives information about the species cluster (based on 95% ANI cut-offs), oral site, and continent. Bootstrap support based on 100 replicates is indicated as percentages for principal branches. The scale bar indicates substitution per position.

Within the *T. forsythia*-group, species cluster 5 corresponds to *T. forsythia,* and there are three species clusters specific to humans (species clusters 3, 4, and 5) (Figure 2a). Species clusters 3 and 5 contain both modern and ancient genomes, while species cluster 4 is composed of a single genome, which has low completeness and high contamination, so this species cluster may be an artifact (Supplementary Figure S4). Species cluster 3 includes the highest proportion of ancient MAGs (53% vs. 16% in species cluster 5), and also contains the oldest sample, which is from Italy and aged 14,800 calibrated years before the present (cal BP)(Ottoni et al. 2021) (Figure 1a, c, 2a). We propose the name Candiatus *Tannerella abscondita* (*T. abscondita* (ab.scon′di.ta. L. fem. adj. *abscondita*, hidden or concealed, referring to the cryptic presence in microbial communities) for species cluster 3. Other species clusters are specific to chimpanzees and gorillas (species cluster 6), brown bears (species cluster 2), or dogs and brown bears (species cluster 1) (Figure 2a), highlighting a deep evolutionary history of oral *Tannerella* in mammals.

In the *T. serpentiformis-*group there are five species clusters specific to humans (clusters 2, 3, 4, 5, and 6), and only species cluster 2 contains an ancient genome, which comes from Germany (900 BP)(Warinner et al. 2014) (Figure 2b, Supplementary Figure S5). This ancient genome was misidentified as *T. forsythia* in 2014(Warinner et al. 2014), and the discovery that it is a different species explains the high SNP counts based on mapping to a *T. forsythia* reference genome. Species cluster 6 corresponds to the uncultivated phylotype *Tannerella* sp. oral taxon 808 (*T*. sp. HMT-808)(Beall et al. 2018), and we propose the name Candidatus *Tannerella ophioides* (Ca. *T. ophiodes*, o.phi.oi′des. Gr. n. ophis, snake; Gr. adj. -oeidēs, resembling; N.L. fem. adj. ophioides, snake-like) for this species cluster. For species cluster 5 we propose the name Candidatus *Tannerella anguiformis* (Ca. *T. anguiformis*, an.gui.i.for′mis. L. anguis, snake or eel; L. adj. -formis, shaped like; N.L. fem. adj. anguiformis, eel-shaped); either Ca. *T. anguiformis* or the unnamed species cluster 2 may correspond to the uncultivated phylotype *Tannerella* sp. HMT-916(Escapa et al. 2018). Other species clusters are specific to chimpanzees (species clusters 7 and 8) or howler monkeys (species cluster 1) (Figure 2a). Overall, we identify three novel human-specific *Tannerella* species clusters, and there appears to be little overlap in *Tannerella* species between humans and other animals. The overall lack of ancient genomes from *T. serpentiformis*-group species clusters suggests that these were historically less prevalent or abundant in dental biofilms than they are currently.

### Niche partitioning in Tannerella species

We next explored niche partitioning on a local scale (within the mouth) and at a global scale (at the continent level) within *Tannerella*, as species or even strain-level niche partitioning has been shown for other oral species including members of the genera *Streptococcus* and *Veillonella,* among others (Mark Welch et al. 2016; McLean et al. 2022; Giacomini et al. 2023, 2025), and this may correlate with distinct genetic content(Shaiber et al. 2020; Velsko & Warinner 2025). To date, all taxa investigated for niche partitioning are all oxygen tolerant early-colonizers, and the extent to which anaerobic, late-colonizer taxa such as *Tannerella* and *Porphyromonas* likewise show niche partitioning has not been extensively investigated.

Within the *T. forsythia*-group, we observed that *T. forsythia* exhibits oral biogeographic partitioning, and is present in both non-shedding (dental plaque, dental calculus) and shedding (saliva) oral habitats (Figure 2b, Supplementary Figure S5). Notably, all MAGs derived from saliva cluster together separately from the MAGs derived from dental plaque and calculus, with the exception of three ancient MAGs, which are also from Oceania (Figure 2b). Ca. *T. abscondita* MAGs were recovered from modern dental plaque as well as both modern and ancient dental calculus, but not in habitats with shedding surfaces (i.e., saliva, tongue, buccal mucosa), suggesting strong oral biogeographic partitioning in this species cluster. The oral niche-partitioning we observe within the *T. forsythia*-group may indicate unique genetic content allowing the bacteria to specialize in colonizing unique oral niches, which we later investigated in a pangenome analysis.

Within the *T. serpentiformis-*group, *T. serpentiformis* shows small regions of the phylogeny with oral biogeographic niche partitioning, as well as some continental geographic specificity (Figure 2c,d), but the oral site and geographic origin of MAGs are not strictly correlated, with MAGs from across the globe found in saliva and on the tongue falling adjacent in the phylogeny. The majority of genomes for this species come from Africa and North America, so poor geographic representation may prevent us from observing stronger phylogeographic signals. However, currently the data suggests that *T. serpentiformis* has higher ecological plasticity than *T. forsythia*. In contrast, Ca. T. *ophioides* and Ca. *T. anguiformis* in the *T. serpentiformis*-group are only found in human dental plaque and calculus, and thus may specialize in biofilm colonization.

### Novel species clarifies Tannerella diversity in ancient dental calculus

Several ancient dental calculus studies have presented evidence of an ancient lineage of *T. forsythia* that lacked a number of known *T. forsythia*-specific virulence factors and has seemingly been replaced globally by the currently recognized *T. forsythia(Honap et al. 2023; Bravo-Lopez et al. 2020; Jackson et al. 2024)*. We noted overlap in samples reported to carry this ancient lineage and those with MAGs in *T. forsythia*-group species Ca. *T. abscondita*, a previously unnamed species cluster, rather than the *T. forsythia* species cluster; we propose that the early *T. forsythia*-group lineage presented in those studies may actually be this novel species. Ancient and modern Ca. *T. abscondita* MAGs are globally distributed, and the species is therefore still globally present. The single modern North American MAG is not clustered with the ancient North American MAGs, perhaps because the individual that MAG came from(Human Microbiome Project Consortium 2012) has a European genetic background and carries a strain with European ancestry.

Investigating the age of the split between species clusters in the *T. forsythia*-group using a BEAST analysis (Supplementary Figure S6), we found a deep split time between these two species, 60,000 ± 5,000 years ago. The dates indicate that speciation of these two organisms is not correlated with medieval population expansion and agricultural intensification(Jackson et al. 2024). Rather, this event occurred at the same time as the later dispersal of humans out of Africa (Sümer et al. 2025) and a similar event was already reported in gut bacterial species(Wibowo et al. 2021). However, the large confidence interval means we interpret the precise split date with caution, and additional genomes assembled in future studies may help narrow this interval.

### Tannerella forsythia*-group pangenome*

We next looked at the pangenome of the *T. forsythia*-group to see if gene content is associated with the phylogeny, oral site, or geographic origins of the MAGs. We saw that nearly a third of genes appear to be core to all species in this group, while slightly more than one third of genes are core to *T. forsythia* (Figure 3a). We did not see any accessory gene content that was restricted to a particular oral habitat, nor did we see any accessory gene content strictly found in or strictly absent in MAGs from Oceania. (Figure 3a). We applied a GWAS approach(Lees et al. 2018) to identify genes enriched in *T. forsythia* MAGs from Oceania compared to elsewhere, and found 10 genes with significant association and high effect size (p < 0.02, effect size > 0.5; Supplementary Figure S7). Among these are the *fur* gene encoding a ferric uptake regulator protein and two genes involved in translation, but the remaining seven are unannotated. We did not investigate the gene association content within the Oceanian clade due to the quality and small number of ancient calculus MAGs (3, Figure 2b). While it is difficult to draw conclusions regarding how gene content distinguishes the Oceanian clade, it is possible that other genomic changes(Mishra & Lercher 2024; Kirsch et al. 2024; Maruyama et al. 2009; Kaul et al. 2026) may be associated with the transition from a predominantly biofilm-associated lifestyle to a predominantly saliva-associated lifestyle.

**Figure 3:**
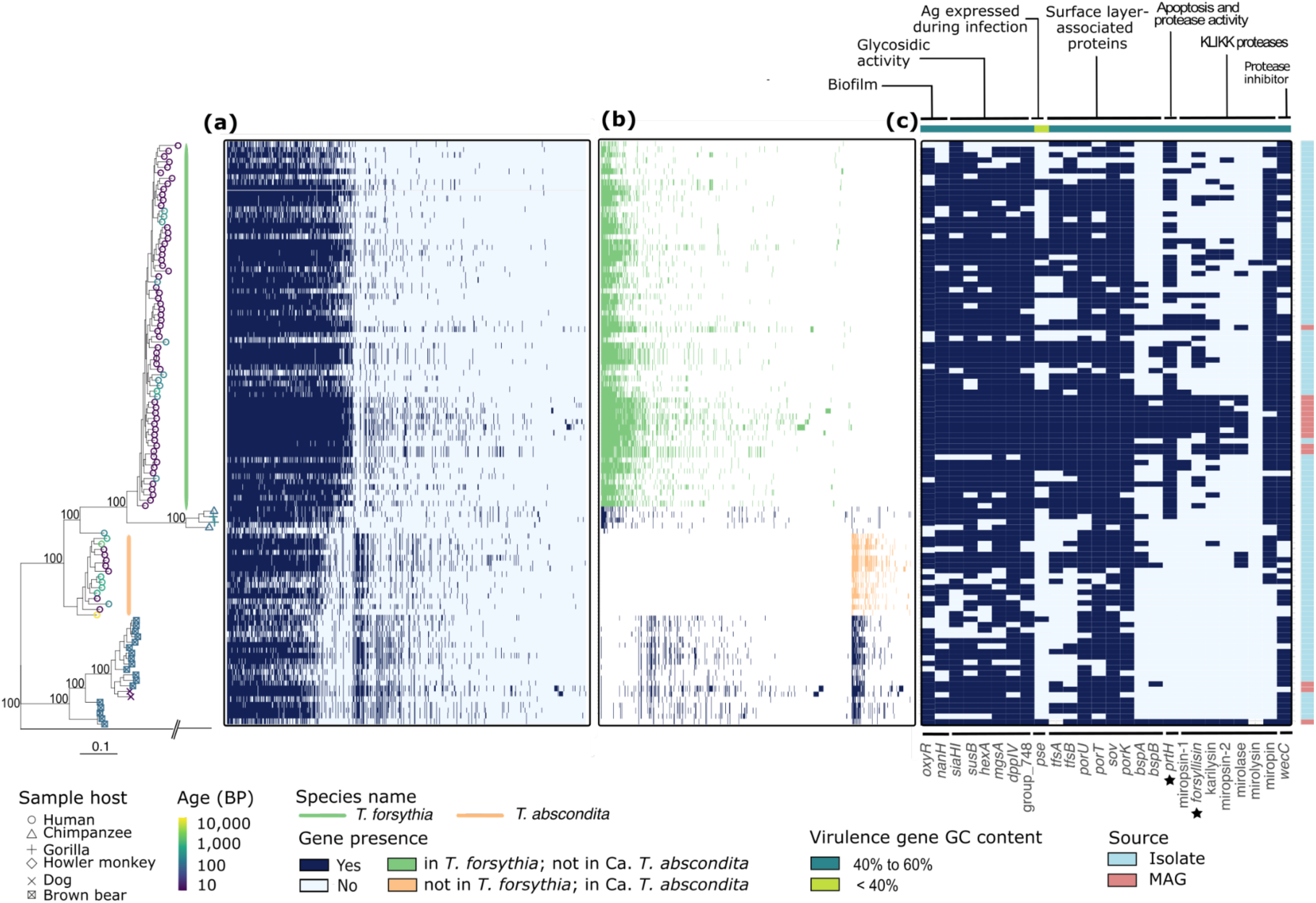
Gene content assessment for the *Tannerella forsythia*-group. **(a)** Phylogenetic tree of *Tannerella forsythia*-group MAGs, as shown in Figure 2a, and pangenome presence-absence matrix. The green vertical bar indicates the *T. forsythia* MAGs, and the orange vertical bar indicates the Ca. *T. abscondita* MAGs. **(b)** Presence-absence gene matrix of genes uniquely present in *T. forsythia* or Ca. *T. abscondita*. **(c)** Virulence factor gene presence-absence matrix. Genes are grouped by activity, indicated across the top of the matrix, along with a colored bar indicating whether the median GC% of each gene is above or below 40%. Stars indicate *prtH* and *forsylisin*. **(d)** Matrix of the genome source.

We next looked at how gene content differs between *T. forsythia* and Ca. *T. abscondita* to better understand the distinctions between them. We found a set of genes (n = 426) that are present in *T. forsythia,* but largely absent in Ca. *T. abscondita*; and a set of genes (n=123) present in Ca. *T. abscondita* that are largely absent from *T. forsythia* (Figure 3b). In both gene sets, the genes are also found in the *Tannerella* species clusters derived from chimpanzees, brown bears, and dogs. The presence and absence of these genes across the phylogeny suggests specific gene loss during evolution of *T. forsythia* and Ca. *T. abscondita*, rather than gene gain. The genes in both species clusters perform a variety of functions, and we were unable to identify any specific metabolic pathways or activities that might differentiate the physiology of *T. forsythia* and Ca. *T. abscondita* through these unique sets of genes, which may be partly due to the high number of unannotated genes (more than 91% for *T. forsythia* and more than 92% for *T. abscondita*) or the incompleteness of the MAGs. Future efforts to isolate and culture Ca. *T. abscondita* are needed to investigate its physiology and biochemistry.

### Tannerella forsythia-group virulence gene distribution

The previously described ancient *T. forsythia* lineage presented in several studies, which we reclassify as Ca. *T. abscondita*, has been reported to lack several key of virulence factors found in present-day *T. forsythia* genomes, prompting speculation that *T. forsythia* acquired these virulence factors relatively recently(Jackson et al. 2024). We therefore explored the virulence gene repertoire of both species to better understand the distribution of these genes and their phylogenetic history (Figure 3c). We found the virulence genes to be well-conserved within the *Hominidae* lineage (Figure 3c), and genes annotated in the functional categories of biofilm, glycolytic activity, and surface layer-associated proteins are shared across all *Tannerella* species (Figure 3c), suggesting they play central roles in the biochemistry of this genus. However, we found a significant enrichment of surface layer-associated virulence factors in primate-specific clades (species clusters 3-6) compared to non-primate clades (species clusters 1 and 2) (chi-square test, p = 9.58×10−6). Because surface layer proteins interact directly with host tissues and metabolites, this may reflect environmental or host specialization.

Three categories of virulence factors appear to be significantly less prevalent in Ca. *T. abscondita* than in *T. forsythia* (chi square test, p = 0.03): surface layer-associated proteins, apoptosis and protease activity proteins, and KLIKK proteases. Among these genes are *tspA* and *tspB*, which form the characteristic S-layer of *T. forsythia* (Figure 3c), *prtH* which can degrade complement component C3, and six genes encoding KLIKK proteases that are secreted and degrade host proteins(Ksiazek et al. 2015): miropsin-1, forsylisin, karylysin, miropsin-2, mirolase, and mirolysin. The genes with the greatest difference in prevalence are *prtH* and forsylisin (indicated by stars in Figure 3c), which are completely absent in Ca. *T. abscondita*, but present in modern and ancient genomes of *T. forsythia*.

The S-layer proteins and KLIKK proteases are exported through a type IX secretion system made up of genes *porU*, *porT*, *sov*, and *porK(Narita et al. 2014)*, all of which appear to be uniformly distributed across MAGs in the *T. forsythia*-group (Figure 3c). The absence of S-layer proteins and KLIKK proteases in Ca. *T. abscondita* therefore does not seem to be due to a lack of machinery to process them. Notably, only the NCBI isolate genomes have a majority of KLIKK proteases consistently detected, suggesting that there may be an assembly bias in reconstructing these genes in MAGs. After visual inspection of the NCBI isolate genomes, we were unable to find evidence that the KLIKK proteases are carried by mobile elements or are in the vicinity of transposes or integrases, although there are Acido-Lenti-1 ncRNAs downstream of miropsin and karylisin. Little is known about the function of Acido-Lenti-1 ncRNAs(Weinberg et al. 2010), but large unannotated stretches between these motifs and the subsequent CDS suggest they may be involved in regulating the expression of these proteases. Overall, it appears that during its evolution *T. forsythia* may have acquired particular virulence factors like as *tfsAB* and the KLIKK proteases, which are largely absent from more deeply diverged *Tannerella* species, but the evolutionary history and origins of these genes remains to be investigated.

### *Genetic diversity in* Porphyromonas gingivalis *and* Porphyromonas pasteri

We next investigated genetic diversity and evolutionary relationships within two characterized species of *Porphyromonas* that are abundant in our MAG dataset: *P. gingvalis* and *P. pasteri*. While we did not recover any MAGs in these species from Neanderthals, we did recover a *Porphyromonas* MAG from a Neandertal in Spain(Fellows Yates et al. 2021) that falls in an unnamed species cluster, primary cluster 11, with two other *Porphyromonas* assemblies from NCBI (Supplementary Figure S2). This suggests that this species has a long history in the *Homo* lineage. In other oral species with both human and Neanderthal representatives, Neanderthal-derived genomes tend to form distinct clades or strains(Velsko et al. 2026), reflecting the deep split times of humans and Neanderthals. However, some strains appear to be shared and may indicate more recent contact. At present, there is insufficient data to resolve the strain structure of *Porphyromonas* species cluster 11, but future recovery of additional MAGs in this cluster may reveal additional information about the evolution of this species and human-Neanderthal interactions.

We found that all *P. gingivalis* genomes form a single primary cluster at 95% ANI (Figure 4a,b, Supplementary Figure S8), which is derived primarily from human hosts and a single chimpanzee. It is present in shedding (saliva, tongue) and non-shedding habitats (dental plaque, calculus) in samples from across the world, including Africa, Europe, Asia, Oceania, North and South America. The majority of *P. gingivalis* genomes in our dataset are from modern, present-day populations, with only six ancient human calculus MAGs recovered. This was not unexpected, given that *P. gingivali*s has been reported as prevalent but not abundant in ancient dental calculus based on metagenomic profiling(Velsko et al. 2019; Velsko & Warinner 2025; Quagliariello et al. 2022), and the relative abundance of this species in dental biofilms is estimated to have increased in the last 200 years(Velsko et al. 2026). The ancient *P. gingivalis* MAGs originate from Rapa Nui (350 BP)(Velsko et al. 2024), Serbia, (8,800 cal BC)(Ottoni et al. 2021), and England (200 BP and 700 BP)(Velsko et al. 2019); (Velsko et al. 2025). We built a maximum likelihood marker gene-based phylogeny to investigate *P. gingivalis* evolutionary relationships in combination with sample metadata (Figure 4a). No phylogenetic patterns were evident for either the sample oral source or for the geographic origin (Figure 4b), which suggests that, as with *T. serpentiformis, P. gingivalis* has a high ecological plasticity. However, the majority of saliva-derived *P. gingivalis* MAGs come from Oceania, suggesting local niche adaptation that may be geographically restricted. Broader sampling and deeper sequencing to obtain additional MAGs from global populations will help to better understand these patterns.

**Figure 4:**
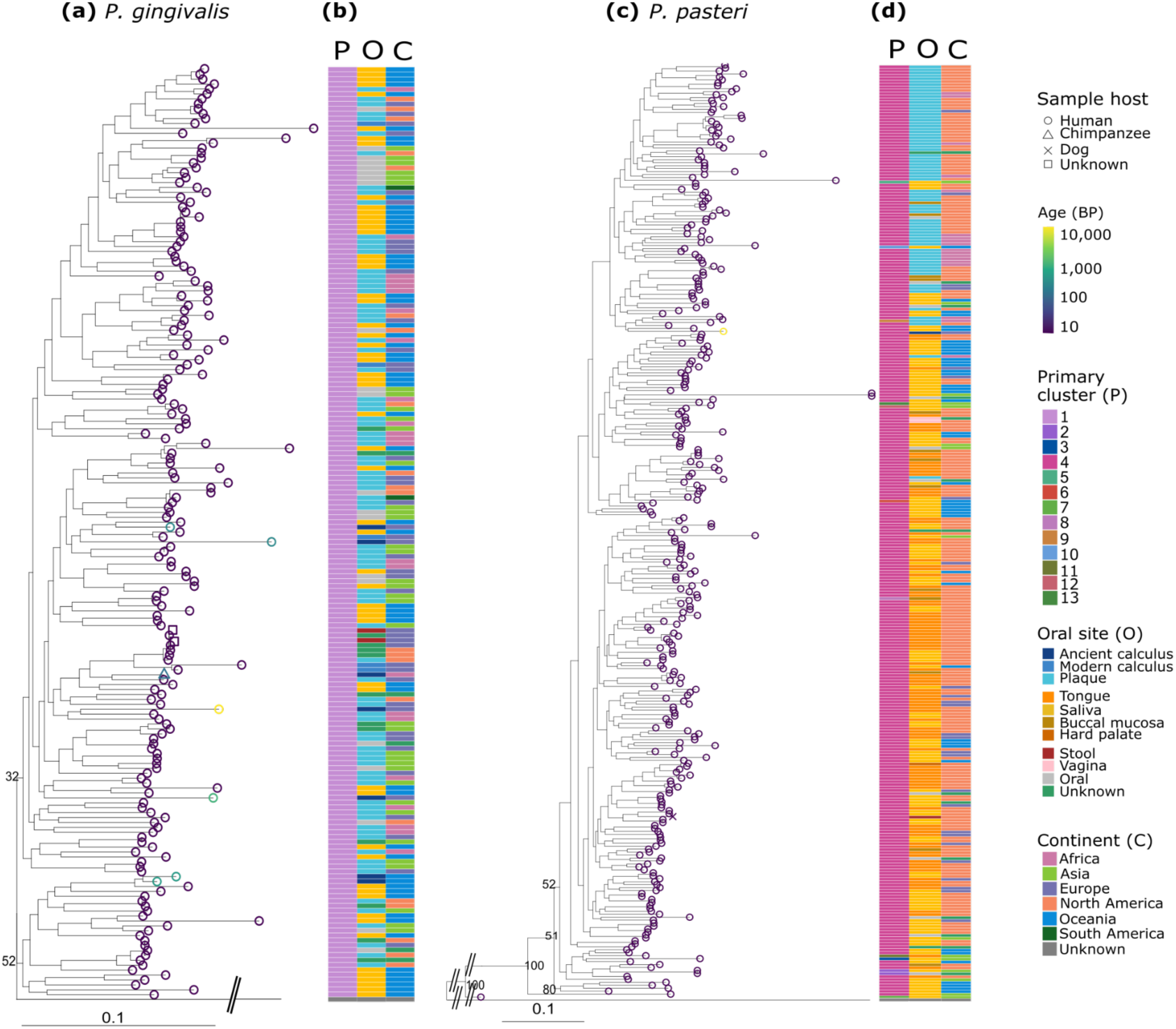
Phylogenetic relationship of *Porphyromonas* species to geographic origin and oral sample site. **(a)** Maximum likelihood phylogenetic tree of *Porphyromonas gingivalis*, rooted on *Porphyromonas pasteri.* **(b)** Metadata matrix of *P. gingivalis*. **(c)** Maximum likelihood phylogenetic tree of *Porphyromonas pasteri*, rooted on *P. gingivalis*. **(d)** Metadata of *P. pasteri*. Tree tip color corresponds to sample age in years before present (BP) and tip shape corresponds to sample host.The matrix gives information about the species cluster (based on 95% ANI cut-offs), oral site, and continent. Bootstrap support based on 100 replicates is indicated as percentages for principal branches. The scale bar indicates substitution per position.

We next looked at genomic diversity in *P. pasteri,* which includes MAGs in multiple 95% ANI species clusters (clusters 2 to 12) (Figure 4c,d, Supplementary Figure S9); however, the inflated number of species clusters represented by single genomes is likely an artifact caused by a combination of lower completeness and higher contamination. Only one MAG is not from a human, and this is from a dog in primary cluster 4 (Figure 4c,d). Like *P. gingivalis*, *P. pasteri* MAGs are globally distributed. While the phylogeny shows a clade with primarily plaque-derived MAGs, nearly two-thirds of the *P. pasteri* MAGs in our dataset come from shedding surfaces (Figure 4d), suggesting that saliva/tongue are the preferred niche of this species. This may partly explain why we identified only a single ancient human calculus MAG in this phylogeny, which is from England, aged 700 years(Velsko et al. 2025).

### P. gingivalis fimA *genotypic diversity*

We next chose to explore genetic variation in the *P. gingivalis* virulence gene *fimA*, which encodes the major fimbriae main structural protein. The major fimbriae mediate bacterial interaction with host tissues(Grover et al. 2014; Amano 2003). Six genotypes of *fimA*, distinguished by multiple single-nucleotide polymorphisms, have been described to date (types I,Ib, II-V), and they have different associations with periodontal disease prevalence(Amano et al. 2000). Type II, along with types Ib and IV, appear to be the most virulent(Nakagawa et al. 2002). Using our *P. gingivalis* MAGs, we investigated the geographic and temporal distribution of *fimA* to better understand its evolutionary history (Figure 5).

**Figure 5:**
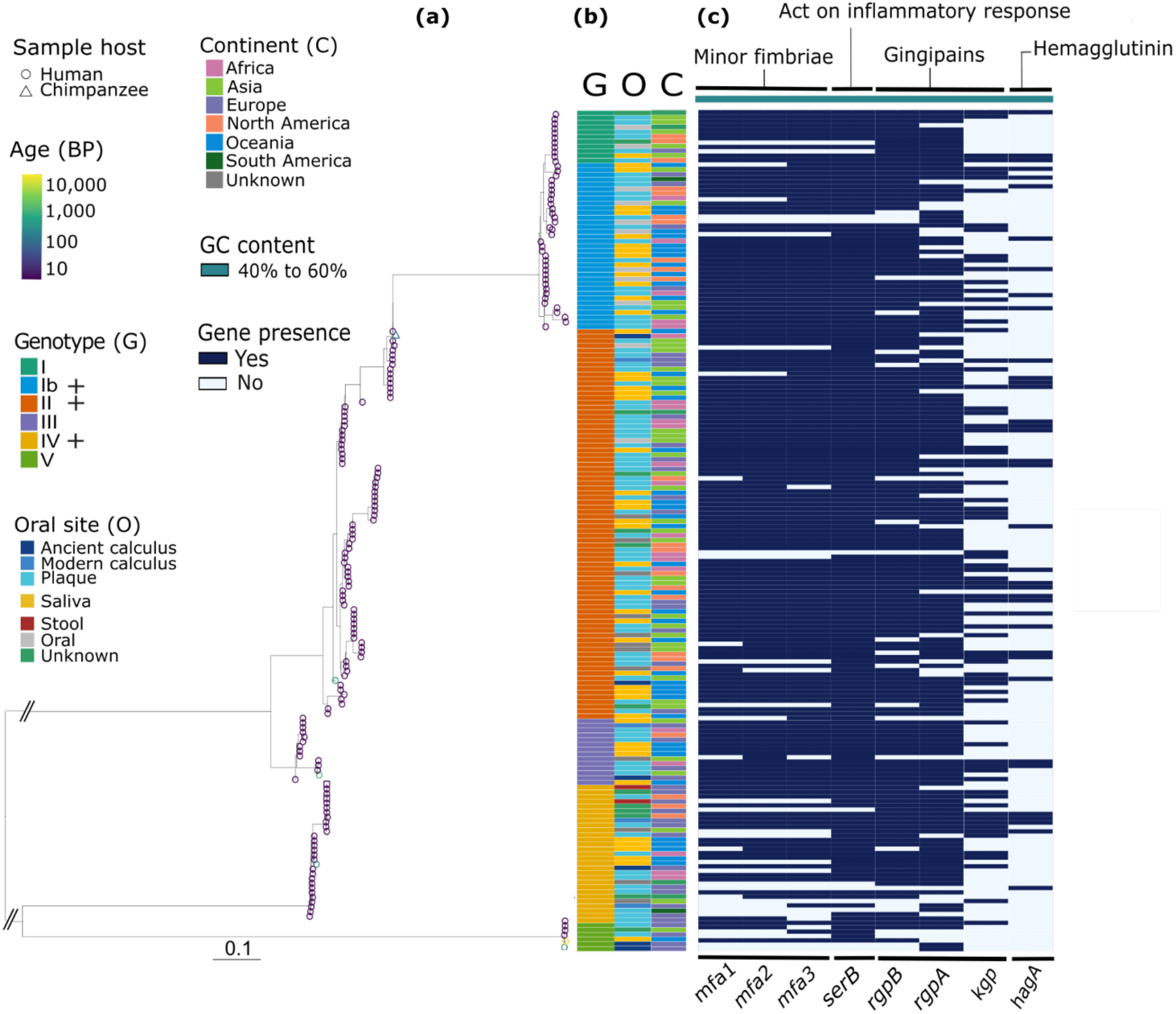
*fimA* genotyping of *P. gingivalis* MAGs and association with prevalence of other known virulence factors. **(a)** Phylogenetic tree of *fimA* gene sequences from *P. gingivalis*. The tree tip color corresponds to the age of the samples in years before present (BP) and shape corresponds to the sample host. The scale bar indicates substitutions per position. A (+) by the genotype indicates that the genotype is strongly associated with periodontal disease. **(b)** Metadata matrix of the *fimA* genotype (G), primary cluster (P), oral site (O), and geographic localization (C) of samples that produced *P. gingivalis* MAGs. **(c)** Virulence factor presence-absence matrix. The function of the genes and the GC content are displayed above the heatmap.

We identified a total of 178 *fimA* genes in 176 MAGs/genomes (93% of the *P. gingivalis* MAGs/genomes), and then extracted the sequences, aligned them, and built a maximum-likelihood phylogeny (Figure 5a). The branching pattern of the tree resembles previously reported phylogenetic clustering of the *fimA* gene based on isolate gene sequences(Enersen et al. 2008), with types I and Ib forming a sister clade to types II and III, while types IV and V are more distantly related. Within each type clade, there are smaller clades with nearly identical sequences suggesting horizontal transfer and diversification across strains; however we do not detect any association with the oral niche or geographic origin of the MAGs (Figure 5b). Ancient samples are distributed in four of the six genotypes: type II (350 BP(Velsko et al. 2024)); type III (700 BP(Velsko et al. 2025)); type IV (200 BP(Velsko et al. 2019)); V (8,800 BP(Ottoni et al. 2021) and 200 BP(Velsko et al. 2019)) (Figure 5a). Among the ancient samples, the individual carrying type IV *fimA* presented some signs of periodontal disease(Velsko et al. 2019); however osteological metadata are not reported for most ancient samples and further inferences cannot be made with the current sample set. We additionally assessed whether virulence factors of *P. gingivalis* co-occur with particular *fimA* genotypes, but no associations were apparent (Figure 5c). With the exception of *kgp* and *hagA*, virulence factors are prevalent across the MAGs, and it is possible that the absence of *kgp* and *hagA* is due to poor assembly because of long nucleotide repeat regions in these genes.

### Limited evidence of genome-encoded antibiotic resistance

We further assessed the presence and distribution of antibiotic resistance genes in *Tannerella* and *Porphyromonas* MAGs to determine whether niche partitioning correlates with resistance profiles, and whether ancient MAGs carry fewer resistance genes. Each resistance gene identified by the Comprehensive Antibiotic Resistance Database (CARD) appears to be broadly distributed among MAGs in the species clusters in which it is identified, and there are no clear associations with MAG age, geographic origin, or oral site. One exception is the mupirocin-like antibiotic resistance gene identified in *P. gingivalis*; however, this hit covered a mean of 4% of the reference gene, and may be a spurious identification Although a variety of genes were identified (Figure 7), the antibiotic resistance repertoire of these species clusters remains unclear, especially considering that rates of resistance reported in clinical isolates of these taxa are low(Kulis et al. 2025; Veloo et al. 2012; Conrads et al. 2021; Andrés et al. 1998).

The most prevalent gene across in the *T. forsythia*-group, *T. serpentiformis*-group, *P. gingivalis*, and *P. pasteuri* MAG sets was “*vanT* gene in *vanG* cluster” (indicated by a star [*] in Figure 7a-d), a gene with alanine racemase activity that is part of a vancomycin resistance gene cassette(Stogios & Savchenko 2020). Panaroo identified the *vanT* gene CDS as the alanine racemase *alr* in the *T. forsythia* and *P. gingivalis* pan genomes, but did not annotate the same gene in *T. serpentiformis* or *P. pasteuri*. We recently showed that vancomycin resistance is the most commonly identified resistance mechanism in oral bacterial genomes(Velsko et al. 2026), and described a highly conserved vancomycin resistance gene cassette found in members of a Gram-positive oral *Anaerovoracaceae* species dating back to the Pleistocene, supporting that this resistance mechanism is ancient(D’Costa et al. 2011).

To determine whether the MAGs in this study likewise carry a full vancomycin resistance cassette, we next examined the surrounding gene architecture. We found a GSCFA-domain containing protein directly upstream of all *vanT* genes, but this two-gene cassette was not part of a longer operon in *Tannerella* or *P. gingivalis* (Figure 7e). The *T. serpentiformis* MAGs carried one of three versions of the gene identified as *vanT*, with differences in gene length, orientation, and completeness. In *P. pasteri*, the gene appears to be the final gene in a six-gene operon that starts with *valS* (a valine-tRNA ligase) and includes an acetyltransferase (GNAT) domain protein CDS (Figure 7f). This operon may be involved in glycopeptide antibiotic resistance, given that GNAT enzymes contribute to this mechanism(Shirmast et al. 2021), and the initial description of *P. pasteri* indicated that the species is resistant to vancomycin(Sakamoto et al. 2015). It seems likely, however, that with the exception of *P. pasteri*, all other genes identified as “*vanT* gene in *vanG* cluster” are strictly alanine racemases rather than the evolutionarily-related *vanT*. Whether the genes identified here are involved in vancomycin resistance remains to be investigated *in vivo*.

### Pseudogene frequency is related to oral site and geography

Although *T. forsythia,* Ca. *T. abscondita,* Ca. *T. ophioides,* Ca. *T. anguiformis,* and *P. pasteri* show oral niche specialization, *T. serpentiformis* and *P. gingivalis* show a flexible adaptability to different ecological niches, suggesting a high genomic plasticity. We observe no significant difference in genome length between non-shedding and shedding surface-derived MAGs (Supplementary Figure S10), however, the presence of pseudogenes and their frequency in genomes from different oral sites may provide insights into this niche adaptation, revealing that functions were dropped as a result of niche specialization(Shaiber et al. 2020). We tested the difference in pseudogene frequency within each species cluster between MAGs from different oral sites (Figure 6); Ca. *T. ophioides* and Ca. *T. anguiformis* were excluded from this analysis because all but one MAG of these species are from a single oral site.

**Figure 6.**
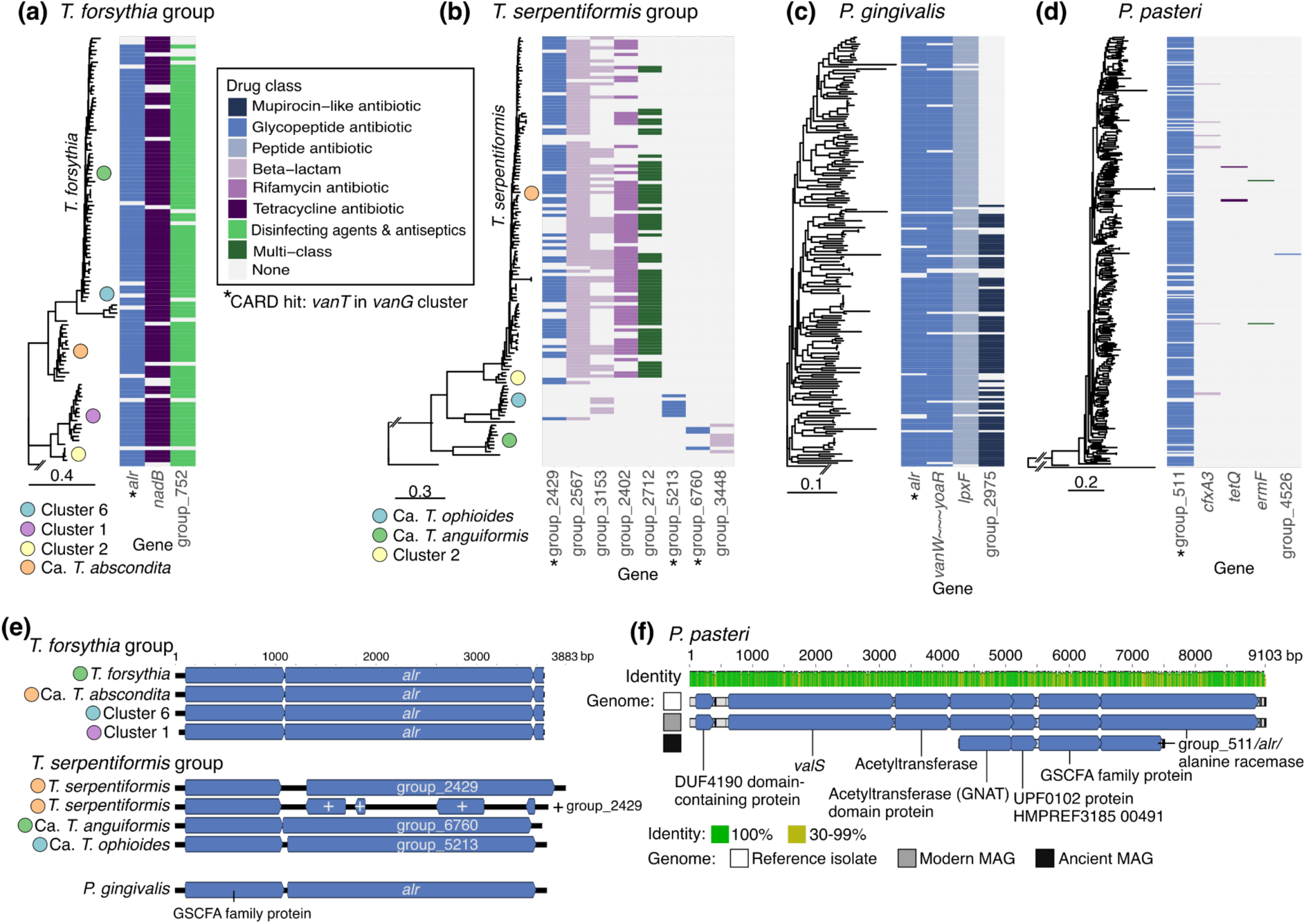
Antibiotic resistance genes in *Tannerella* and *Porphyromonas* species. **A-D**. Gene presence/absence matrices aligned with phylogenetic marker-gene trees. Genes are colored based on the drug class that the resistance mechanism targets. Gene cluster names come from the Panaroo table. **(a)** *T. forsythia*-group. **(b)** *T. serpentiformi*s-group. **(c)** *P. gingivalis*. **(d)** *P. pasteri*. **(e)** Representative examples of the “*vanT* in *vanG* cluster” hit (genes indicated by * in panels **a-d**) identified by the CARD database in selected genomes from the *T. forsythia-*group, the *T. serpentiformis-*group, and *P. gingivalis*. The GSCFA family protein directly upstream is also shown. **(f)** Examples of the *P. pasteri* gene architecture of the operon containing the *alr* gene identified as “*vanT* in *vanG* cluster” from an isolate genome (GCA_001553165), a present-day MAG from Cameroon (CMC001.bin.r.55), and an ancient MAG from Germany (ERSX0002001-megahit_028). Gene names are taken from Bakta annotations because Panaroo did not assign names (i.e., all were “group_####”).

**Figure 7:**
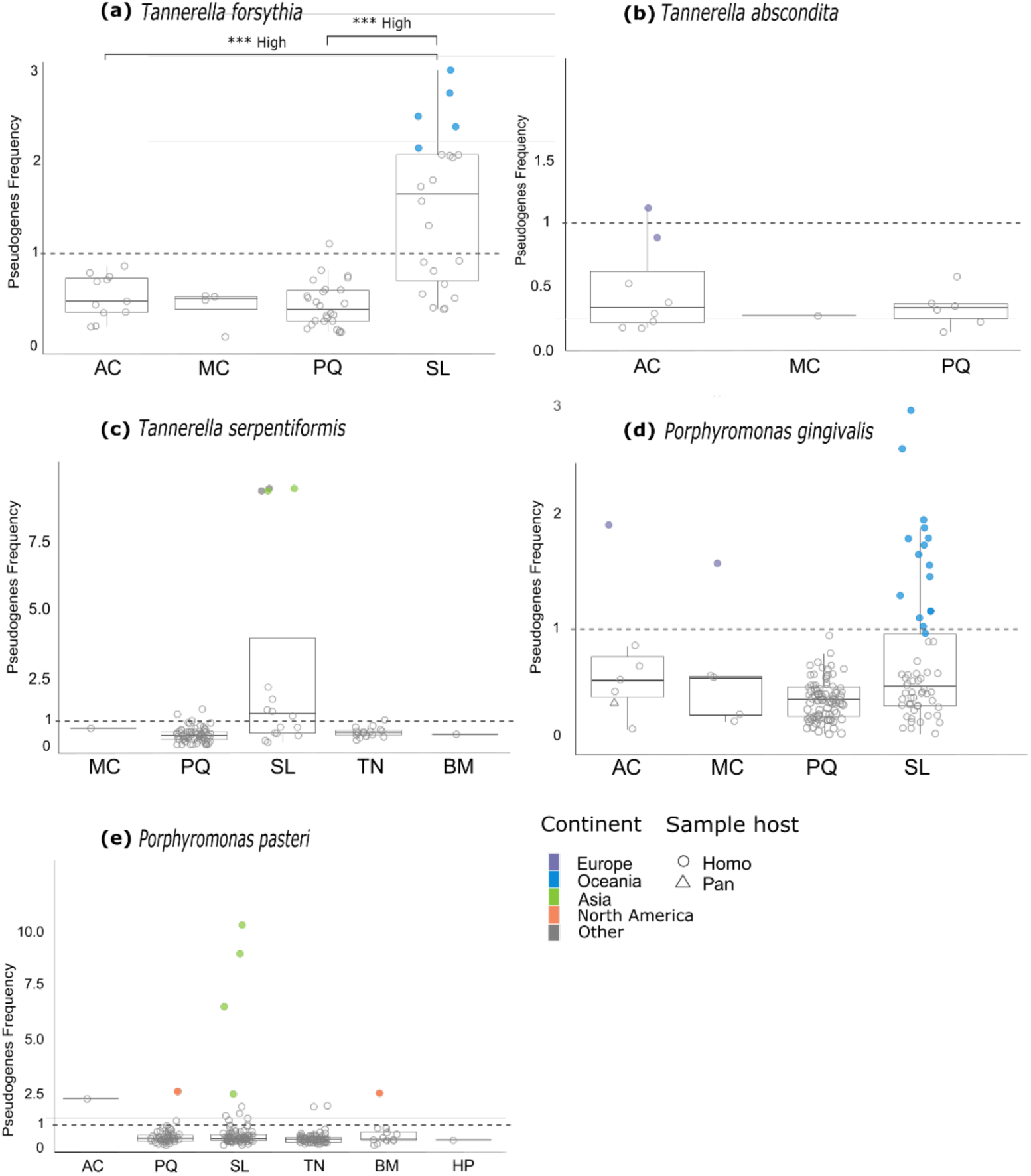
Pseudogene frequency is higher in Oceanian and Asian MAGs from saliva. Boxplots of the pseudogene frequency in MAGs/genomes of **(a)** *Tannerella forsythia*, **(b)** Ca. *Tannerella abscondita*, **(c)** *Tannerella serpentiformis*, **(d)** *Porphyromonas gingivalis*, and **(e)** *Porphyromonas pasteri*, separated by oral site. The geographic origin of the sample is indicated in color only for outliers, defined as points above the highest boxplot upper third quartile across all oral sites within a species. *** p < 0.001; effect size High indicates an effect size > 0.500. A dotted line at pseudogene frequency of 1 is included in all charts to improve comparisons across figures due to the different y-axis scale in each.

We found that pseudogene frequency is higher in MAGs derived from saliva samples than all other oral sources across all species. For *T. forsythia,* this difference is significant (p < 0.05, effect size > 0.5) (Figure 6a). In addition, all the outliers for saliva-derived *T. forsythia* and *P. gingivalis* MAGs are from Oceanian samples, and from Asian or North American samples for *P. pasteri* and *T. serpentiformis.* The majority of these pseudogenes were unannotated, and therefore we cannot determine which functionalities are being lost. Sampling of salivary metagenomes from additional global populations is needed to clarify whether the pseudogenization we observe is restricted to the species investigated here, or a general trend in oral species, as well as whether this pseudogenization is geographically restricted.

## Discussion

Within the oral microbiome, the genera *Tannerella* and *Porphyromonas* are best known for two species strongly associated with periodontal disease, *T. forsythia* and *P. gingivalis*. Although more diversity is recorded in both genera, few isolates and reference genomes of these species exist, and little is known about their evolutionary history or role in the oral microbiomes. Using *de novo* assembly of oral metagenomes from multiple shedding and non-shedding oral habitats, including ancient dental calculus, we recovered a large number of genomes representing unexplored diversity in both *Tannerella* and *Porphyromonas*, and expanded representation of documented species. Using this large dataset we identified Ca. *T. abscondita*, a new *Tannerella* species that was previously mis-identified as *T. forsythia* in ancient metagenomes, and the new species Ca. *T. ophioides* and Ca. *T. anguiformis* that fall within the broader *T. serpentiformis*-group. We observe differences in niche specialization and geographic partitioning across *Tannerella* and *Porphyromonas* species, which in some cases is significantly associated with pseudogenization in saliva. Our approach reveals insights into oral microbial niche tropism and ecology at an unprecedented global and temporal scale.

### Cryptic species diversity in metagenomes can be revealed through cautious selection of reference genomes

One of the most notable findings is the discovery of novel species diversity within human *Tannerella*, including three new well-supported species clusters, for which we propose the names Candidatus *Tannerella abscondita*, Candidatus *Tannerella ophiodes*, and Candidatus *Tannerella anguiformis*. Ca. *T. abscondita*, which is closely related to *T. forsythia*, is particularly noteworthy because it was previously described as an unusual avirulent lineage of *T. forsythia* in ancient human dental calculus. However, based on ANI, phylogenetic clustering, and gene content, Ca. *T. abscondita* appears well-supported as an independent species. Within Ca. *T. abscondita* there may be continent-specific lineages, but additional genomes are needed to clarify the phylogeographic structure, and additional sampling of modern-day populations, particularly those in North America, is needed to determine if the species is no longer present in indigenous populations, as has been suggested(Bravo-Lopez et al. 2020; Honap et al. 2023; Blevins et al. 2026).

After separating the two species, we do not find evidence that *T. forsythia* has recently acquired virulence genes as has been suggested(Jackson et al. 2024), but rather that the gene content of Ca. *T. abscondita* and *T. forsythia* differ, including the presence of particular virulence genes. The Ca. *T. abscondita* MAGs were recovered from samples both with and without evidence of periodontal disease. Given that both *T. forsythia* and *T. serpentiformis* are associated with periodontal disease(Ansbro et al. 2020), it is possible that the genus *Tannerella* is primed to take advantage of an inflammatory environment. Some as-yet-undescribed virulence factors present in Ca. *T. abscondita* may be implicated in pathogenicity in future *in vitro* studies, and it is likely the species carries a distinct virulence gene repertoire.

The mistaken identification of Ca. *T. abscondita* as a lineage of *T. forsythia* likely stems from the mapping-based approach used in its original discovery, wherein a reference genome, in this case *T. forsythi*a, was preselected and assumed to correctly represent diversity in the samples. As mentioned in the results, we mis-identified the most abundant *Tannerella* species in 900 year-old calculus from a nun in Germany as *T. forsythia* using a reference-based mapping approach, while the MAG recovered from this sample is *T. serpentiformis*. Similarly, we identified *T. forsythia* in howler monkey ancient dental calculus through mapping to a reference genome(Fellows Yates et al. 2021); however, ANI and phylogenetic analysis of the *Tannerella* MAGs recovered from these samples clearly indicate that they belong to an independent species more closely related to *T. serpentiformis*. A reference mapping-based approach to recover ancient microbial genomes works well in studies of well-described pathogenic microbes that are genetically distinct from microbial commensals and environmental contamination, such as *Yersinia pestis*, the causative agent of bubonic plague(Swali et al. 2023). However, in complex microbial communities such as dental biofilms, greater caution is warranted to ensure that an accurate reference genome is chosen. To this end, recent large-scale MAG recovery studies provide novel reference genomes that substantially expand the diversity of oral species currently represented in databases such as NCBI GenBank(Velsko et al. 2026; Cha et al. 2025), and offer a more comprehensive collection of references with which to explore (ancient) oral microbiome diversity.

### Oral niche adaptation and geographic restriction

We observe distinctive oral biogeographic and geographic restrictions across the species we investigated. Representation of *Tannerella* in samples from Asia was extremely low, consisting of only four *T. serpentiformis* MAGs, however it is unclear whether this is an artefact of selection bias in the MAGs made publicly available for the largest Asian study(Zhu et al. 2021) we included.The majority of *Tannerella* MAGs come from non-shedding biofilm samples (dental plaque, calculus), while MAGs from saliva and tongue tend to cluster together in the phylogenies. This pattern suggests that the primary niche of this genus is dental biofilms, but particular strains have undergone niche transitions to adapt to shedding habitats, with *T. serpentiformis* showing the highest ecological plasticity in the *Tannerella* genus. In contrast, *Porphyromonas* shows a different pattern. The majority of *Porphyromonas pasteri* MAGs are derived from saliva and tongue samples, indicating that the primary niche for this species is shedding habitats. However, like the saliva-derived MAGs from *T. forsythia*, the majority of plaque-derived *P. pasteri* MAGs fall within a single clade, suggesting a single transition event and subsequent diversification. *P. gingivalis* shows more ecological plasticity than *P. pasteri*, which is unexpected given that *P. gingivalis* is strongly associated with subgingival plaque and periodontal disease. This finding suggests that gene content variation within this species allows for diverse niche adaptation.

In contrast to *Tannerella*, MAGs of both *P. gingivalis* and *P. pasteri* were recovered globally, and no phylogeographic signal is evident in either species. Unexpectedly, few *Porphyromonas* MAGs were recovered from chimpanzees, despite the high relative abundance of this genus based on metagenome taxonomic profiling(Fellows Yates et al. 2021). One possible explanation is that high species diversity in these samples interfered with binning algorithms(Velsko et al. 2026), although further work is needed to investigate this. In *P. gingivalis,* recombination events may be frequent(Frandsen et al. 2001), which correlates with the lack of phylogeographic or oral biogeographic signals in our dataset. In addition, *P. gingivalis* carries a diverse repertoire of prophages(Matrishin et al. 2023), and while we did not investigate the diversity of prophages in our MAGs, differences in phage carriage and lytic potential may contribute to the signals we observe. Given the clear oral biogeographic signal in *P. pasteri,* whether this species experiences similar levels of recombination or phage lysogeny is of interest to investigate.

Although *P. gingivalis* is primarily considered to inhabit subgingival habitats, we identified a number of MAGs of this species from saliva samples. Given that *P. gingivalis* can be found at the surface of dental biofilm structures(Zijnge et al. 2010), and is relatively aerotolerant(Nakayama 1994; Xie & Zheng 2012; Romero-Lastra et al. 2017), this finding is perhaps not surprising. *P. pasteri*, however, appears to be primarily an inhabitant of shedding surfaces that later experienced a transition to a biofilm lifestyle. In dental biofilms, corncob structures sporting *Porphyromonas* cells are predicted to carry *P. pasteri*(Morillo-Lopez et al. 2022), so it is of interest to investigate whether the biofilm-derived *P. pasteri* genomes contain evidence of metabolic interdependency with the *Corynebacterium* and *Streptoccocus* species that make up the structural center of corncob structures.

In Oceania-derived MAGs, pseudogenization and iron uptake regulation appear to be a significant part of genomic adaptation from a dental biofilm niche to saliva. One possible alternative explanation is that saliva-derived bacteria may be residing intracellularly in epithelial cells(Mishima & Sharma 2011; Colombo et al. 2007) that were shed into saliva and were released from the host cells during sample processing and DNA extraction. In this case, pseudogenization may be a result of adapting to intracellular living. The small cluster of ancient dental calculus Oceania-derived *T. forsythia* MAGs that fall together with modern-day saliva MAGs from Oceania in the *T. forsythia*-group phylogeny suggests that some genomic evolution occurred in *Tannerella* species in Oceania prior to the niche transition. We also observed that the *P. gingivalis* saliva-derived MAGs are nearly all from Oceania, and the two ancient calculus-derived MAGs from Oceania fall in a cluster with several Oceanian saliva samples, again suggesting a geographically localized oral niche transition. Oceanian populations are known for consuming kava (*Piper methysticum*), a plant that contains anti-inflammatory compounds that have been shown to reduce gingivitis(Huck et al. 2018; Cai et al. 2018), and may thus impact microbial colonization of gingival tissues. Given that human colonization of Oceania is a relatively recent event in human history(Matisoo-Smith & Gosling 2025), further investigation of oral microbes from Oceanian populations may reveal how quickly host-associated microbes adapt to changes in host behavior(Velsko et al. 2024), and which forces strongly drive host-associated microbial evolution.

### Virulence factor detection and distribution

Virulence factors from *T. forsythia* and *P. gingivalis* appear to be globally distributed, and found in populations with differing degrees of industrialization and urbanization, but the detection of some virulence genes is challenging. In MAGs where virulence genes are missing, this may be due to incomplete assemblies related to limitations of the assembly process, such as biases due to GC content and nucleotide repeat sequences. For instance, the *pse* gene in the *T. forsythia*-group, which synthesizes pseudaminic acid, has a low GC content (35.21%), and may be impacted by the loss of AT-rich DNA fragments in ancient DNA(Mann et al. 2018; Fagernäs et al. 2020), such that there were not enough reads present to assemble the gene (Figure 4c). Loss of AT-rich sequences will also affect mapping-based efforts to reconstruct ancient genomes, and therefore gene GC content should be considered when genes are poorly reconstructed. The surprising lack of the surface-associated leucine-rich repeat genes *bspA* and *bspB* across our *Tannerella* genomes may be due to repeat regions found within these genes, which graph-based assembly programs struggle to reconstruct. For gene-centric studies, reasons for gene non-detection may need to be investigated on an individual basis through additional means such as mapping of short reads to reference genes of interest.

Within *P. gingivalis*, nearly identical sequences of *fimA* are found on different continents, supporting that the species is readily shared between individuals and not geographically restricted, or that horizontal transfer is frequent in this species. The presence of *fimA* in a MAG from a wild chimpanzee ancient calculus sample suggests that the gene has a deep evolutionary history in *Porphyromonas*. Genotypes I and Ib fall on the most shallow tree branch, and may be a more recent development, perhaps related to the observed rise in relative abundance of *P. gingivalis* in dental biofilms within the last 200 years(Velsko et al. 2026). It is of interest to screen additional ancient calculus metagenomes to confirm whether genotypes I and Ib are indeed absent from genomes prior to 200 years ago.

Overall, we find that the red complex bacteria *T. forsythia* and *P. gingivalis*, as well as their newly discovered relatives Ca. *T. abscondita*, Ca. *T. ophioides*, and Ca. *T. anguiformis*, show distinct ecological niche adaptations and evolutionary trajectories, highlighting that although these species live in complex communities with many other taxa, they behave in an individual manner and need to be studied individually. In cases where species are difficult to culture or rare to detect in oral microbial samples, and therefore have few if any isolate genomes for study, metagenome assembly is a valuable approach to reconstruct and represent diverse species. Phylogenetic and pangenome analyses of MAGs from diverse sources can reveal ecological and evolutionary adaptation that deepen our insight into how commensal microbes interact and evolve with their hosts.

## Methods

### Data acquisition

Metagenome assembled genomes (MAGs) taxonomically identified as members of the genera *Tannerella* and *Porphyromonas* from oral sources were subset from a large collection of modern and ancient MAGs(Velsko et al. 2026) (Supplemental Table S1). We selected MAGs with MIMAG-defined (Bowers et al. 2017) high quality (>= 90% completeness and < 5% contamination) or medium quality (>=50% completeness and <10% contamination) determined by checkM(Parks et al. 2015) for further analysis. This includes MAGs from different hosts and different oral sites: humans from saliva, buccal mucosa, tongue, keratinized gingiva, hard palate, dental plaque, and dental calculus); non-human primates including howler monkeys (*Alouatta*), gorillas (*Gorilla*), chimpanzees (*Pan*); and bears (*Ursus*), all from ancient dental calculus. Samples are categorized as coming from shedding (saliva, buccal mucosa, tongue, keratinized gingiva, and hard palate) and non-shedding oral surfaces (plaque and dental calculus) as well as non-oral host-associated sources (vaginal, stool, and gut), across different time periods (from 14.800 years before present (BP) to modern times) and along six continents (North America, Europe, Africa, Oceania, South America, and Asia) (Figure 1).

We added reference genomes taxonomically classified as members of the genera *Tannerella* or *Porphyromonas.* They were identified in the GTDB version r214(Parks et al. 2022) (Supplemental Table S2) and downloaded from NCBI using Entrez database(Sayers 2022). Overall we have 238 genomes for *Tannerella* (48 ancient dental calculus MAGs, 162 modern MAGs (7 calculus, 39 saliva, 17 tongue, 1 buccal mucosa and 98 plaque), and 28 reference genomes (18 plaque, 2 saliva, and 8 unspecified sites). For *Porphyromonas* we have 976 *Porphyromonas* genomes, 18 ancient dental calculus MAGs, 763 modern MAGs (14 calculus, 265 saliva, 347 plaque, 25 buccal mucosa, 6 keratinized gingiva, 1 hard palate, and 105 tongue), and 195 reference (28 saliva, 56 plaque, 69 unspecified oral, 1 gut, 2 stool, 3 skin, 2 vagina, and 34 from unspecified sites).

### Species clustering and selection

To determine the number of species present in our dataset for each genus, we clustered the genomes by their average nucleotide identity (ANI) using dRep version 3.4.3(Olm et al. 2017) with a cut-off of 95% ANI to define species clusters (classical threshold to define species in prokaryote (Konstantinidis & Tiedje 2005)). We treated each 95% ANI cluster as an individual species. To make figures and cluster numbers easier to refer to, we re-numbered the clusters for figures starting each group from cluster 1. Supplemental table S1, S2 lists the original dRep cluster number and the new number used in the figures for cross-reference.

For *Tannerella*, species clusters had clearly-defined boundaries based on ANI, they clustered into two large groups with a mean ANI of 65% between them. To explore the genomic diversity and phylogenetic relationships of *Tannerella* species, we used all *Tannerella* genomes and worked with the two 65% ANI clusters independently. Each of the two *Tannerella* groups included a named species, *T. forsy*thia, and *T. serpentiformis,* respectively, and we refer to the larger 65% ANI group, which includes *T. forsythia*, as the *T. forsythia* group and to the smaller 65% ANI group, which includes *T. serpentiformis,* as the *T. serpentiformis*-group. This left us with 107 MAGs in the *T. forsythia-*group (from humans: 20 ancient calculus, 5 modern calculus, 32 plaque, and 22 saliva, 4 site-unspecified MAGs; 2 gorilla calculus, 2 chimpanzee calculus, 2 dog, and 18 brown bear calculus) and 131 MAGs in *T. serpentiformis*-group (from humans: 1 buccal mucosa, 84 plaque, 17 tongues, 19 saliva, 1 ancient calculus and 2 modern calculus, and 2 unspecified sources, 2 chimpanzee calculus and 3 howler monkey calculus).

In *Porphyromonas,* because of the high genomic diversity in this genus, we looked for species clusters with at least 100 genomes and at least one MAG from an ancient dental calculus source on which to perform further genomic and phylogenetic analyses. We selected the two species clusters meeting these criteria: one included genomes taxonomically classified as *P. gingivalis*, and the other included genomes taxonomically classified as *P. pasteri* (Supplemental Figure S2). An ANI cut-off between 93%-95% is recommended for multiple oral genera to prevent artificially splitting species into multiple clusters(Velsko et al. 2026), and for the *P. pasteri* cluster, we added 11 clusters with 12 genomes in total to the main *P. pasteri* cluster as the ANI was very close to 95% (between 93,95% and 94.95%) (Supplement figure S2). This included 315 MAGs for *P. pasteri* (from humans: 118 saliva, 94 tongue, 13 buccal mucosa, 1 ancient calculus, 72 plaque, 11 oral, 2 vagina, and 3 unspecified samples source; 1 dog stool sample) and 189 MAGs for *P. gingivalis* (from humans: 57 saliva, 84 plaque, 11 calculus, 21 oral samples, and 3 unspecified sites; 1 chimpanzee calculus sample and 2 environmental samples).

### Core and universal gene phylogeny construction

We used a marker gene construction approach to build a phylogeny of each selected cluster using Phylophlan(Segata et al. 2013) version 3.1 and a database of marker gene proteins from *Tannerella forsythia* for both *Tannerella* cluster phylogenies or *Porphyromonas gingivalis* for both *Porphyromonas* phylogenies. Phylophlan was run on medium diversity mode with the following settings: DIAMOND(Buchfink et al. 2015) was used for creating and indexing the database of genes for the output genomes, and MAFFT(Katoh & Standley 2013) was run for the multiple-sequence alignment, trimAl(Capella-Gutiérrez et al. 2009) was used to perform the trimming of gappy regions to filter portions of the alignment that contain a high number of gaps. For inferring phylogeny, FastTree(Price et al. 2009) was used to build the first tree, and raxml(Stamatakis 2014) to build the final tree. We added the ‘--accurat’ and ‘-mutation_rates’ flags to provide a more accurate phylogenetic reconstruction.

The outgroup for the *T. forsythia-*group tree is *T. serpentiformis (*GCF_003033925) and for the *T. serpentiformis*-group is a MAG derived from the howler monkeys (i.e., ERS3774378-megahit_009), as the ANI of this group of genomes is more distant (75%) than for any other species clusters in this group. We rooted the *P. gingivalis* tree on the outgroup *P. pasteri* (GCA_943914445), while we used a midpoint root for the *P. pasteri* tree, which contained the outgroup *P. gingivalis (*GCF_000467855). Bootstraps were performed independently of Phylophlan using RAxML-NG(Kozlov et al. 2019) with identical settings to RAxML within Phylophlan, with 100 replicates. Matrices were used to visualize the sample oral origin, host, age, and geographical location in relation to the phylogeny. We also examined completeness and contamination in relation to the trees (Supplementary Figures S4, S5, S8, S9) to make sure phylogenetic clustering was not due to a lack of completeness or high contamination. Ages were log10-transformed for better visualization in the plots. All trees and heatmaps were generated in R with ggtree(Yu 2020) and ggpl(Wickham 2016) unless otherwise noted.

To confirm the phylogeny structure of the *T. forsythia*-groupand *T. serpentiformis*-group trees we used the 400 universal marker genes from the Phylophlan database(Segata et al. 2013), and generated trees in an identical manner as above. This allowed us to confirm that the phylogenetic relationships of the species clusters was not the result of using marker genes from a specific species.

### Genome annotation and pangenome analysis

To annotate genomes, we used Bakta(Schwengers et al. 2021) v1.9.2, in meta mode skipping CRISPR array detection and with the full database. To determine the pan-genome we used Panaroo(Tonkin-Hill et al. 2020) version 1.5.0. We used the flags ‘--clean_mode strict’ and ‘--remove-invalid-genes’ to avoid contamination or invalid annotations and set the aligner to use MAFFT(Katoh & Standley 2013). We assessed whether the universal annotation settings in Bakta were sufficient for both genera by further annotating all genomes/MAGs with Bakta using a species-specific training file from Prokka(Seemann 2014). The genome on which to base the training was selected from all genomes included in the study as the one with the fewest contigs, highest completeness, and lowest contamination for each species, respectively (Supplementary Table S5). Unless otherwise stated, the untrained annotations were used for functional assessments as more annotations per genome were generated with this approach.

### Virulence factor assessment

We identified virulence-related genes in our MAGs/genomes using the Panaroo gene cluster matrices. Virulence factors were selected from published lists of virulence factors in *T. forsyt*hia(Fellows Yates et al. 2021; Philips et al. 2020; Ksiazek et al. 2015; Narita et al. 2014) and *P. gingivalis(Fellows Yates et al. 2021)* (Supplementary Table S3). For *Tannerella,* to identify the virulence factors based on locus tags listed in Philips, et al.(Philips et al. 2020) we added the reference genome (GCF_000238215) containing the locus tags of interest (Supplementary Table S3) to Panaroo for clustering. Panaroo was run by including 2 versions of all genomes - one version annotated without training, and one annotated with training (see section Genome annotation and pangenome analysis for an explanation of training), in addition to the NCBI annotated reference for *Tannerella forsythia*. All gene clusters in *T. forsythia* representing the virulence factors of interest were identified as those clusters containing the reference genome locus tags. For *P. gingivalis*, all virulence factors were identified by Panaroo gene cluster annotation. To ensure we identified all virulence factors, we first pulled the presence of the virulence factor genes from the untrained genomes, then for genomes where no virulence factor was identified, we searched for the gene in the trained genome. If found in the trained genome, it was counted as present. To assess if the presence of virulence factors between clusters in the *T. forsythia-*group was not random we performed a chi-square test(Yekutieli & Benjamini 1999) using rstatix(Kassambara 2020). We also measured the GC content of genes, using the GC calculator (<u>GC Content Calculator - Online Analysis and Plot Tool - BiologicsCorp</u>), to check if the lack of detection was due to a low GC content. AT-rich sequences appear to be more readily lost from ancient DNA, potentially biasing the assembled low GC content genes (Supplementary Table S3).

### fimA phylogeny

We generated a phylogeny of the *Porphyromonas gingivalis* virulence factor gene *fimA* (fimbrillin), a structural protein of the long or major FimA fimbriae. There are 5 genotypes of *fimA*, which were shown to modulate the virulence of the strain carrying each genotype(Nakagawa et al. 2002). We extracted both the *fimA* and group_2895 gene presences from the Panaroo output gene matrix as they are both designated *fimA* in the annotation column of the Panaroo output table. We corrected the presence absence matrix as described in the *Virulence factor assessment* section. To determine the genotype of each gene sequence, we performed a BLAST search for each gene identified in our MAGs/genomes against reference sequences with an associated genotype(Nagano et al. 2013) (Supplementary Table S4). Genes were required to have a 95% identity to one of the reference genes in the BLAST search to call a genotype. Only a single genome had both *fimA* and group_2895, yet both sequences were identified as type II. In two MAGs, two partial sequences on different contigs, corresponding to different regions of the gene, were identified as *fimA*; for both MAGs, each sequence was assigned the same genotype. We concatenated the two gene alignment files from Panaroo (*fimA*, group_2895) into a single fasta file, re-aligned the sequences using MAFFT, built a tree using RAxML with the GTR+G model, and built and bootstrapped a maximum-likelihood tree using raxml-ng(Kozlov et al. 2019) with 100 replicates.

### Identification of pseudogenes

Bakta annotates genes as pseudogenes if it detects indels, mutations, or an early stop codon, so we used the Bakta output from the untrained annotation file from Prokka. to determine the number of pseudogenes in our dataset. We looked at the frequency of pseudogenes (determined as the proportion of pseudogenes out of total genes) in MAGs/genomes from each oral site. Statistically significant differences in the pseudogene frequency between oral sites were determined with a pairwise Wilcoxon test and Benjamini-Hochberg multiple test correction. We also calculated the effect size to determine the magnitude of the relationship between our variables (Figure 6). We assessed whether there was a trend between the geographic origin of the sample and pseudogene frequency by looking at the geographic origin of outliers, defined as MAGs/genomes above the highest boxplot third quartile across all oral sites within a species.

We also looked for a difference in the size of the genomes between shedding and non-shedding surfaces by multiplying the length of each MAG in bp by the length of the reference genome, GCF_000238215 for *Tannerella* (3405521 bp and 98.91% complete), and GCF_000010505 for *Porphyromonas* (2354886 bp and 99.92% complete) and dividing by the completeness of the MAGs (Supplementary Figure S10). Finally, we investigated the completeness and contamination of outliers to make sure that these samples did not have lower completeness or higher contamination that might lead to a higher estimated pseudogene count (Supplementary Table S1, S2). Boxplots were generated using ggplot2(Wickham 2016) and statistics were calculated using rstatix(Kassambara 2020).

### Beast analysis

To estimate the age of the splitting of the two human-specific species from the *T. forsythia*-group, we looked at the core genome alignment obtained from Panaroo using the untrained annotation file from Prokka. We defined the core genome as genes shared across 95% of the genomes. Thus, the alignment contains 13 core genes and 3,959 single nucleotide polymorphism (SNPs). BEAST v2.7(Bouckaert et al. 2019) was run with a GTR-γ4 substitution model. We used a strict clock model as it has been shown that using different clocks does not improve the result for other bacteria(Wibowo et al. 2021) and a coalescent Bayesian skyline, as it is the most suitable model for other bacteria(Wibowo et al. 2021; Jackson et al. 2024). We first ran BEAST following preliminary run recommendations: a Markov chain length of 10^6 with a 10% burn-in. We then ran BEAST with a chain length of 9×10^7 to have an effective sample size (ESS) > 100. This is to make sure to have a good convergence and reliability of the Markov Chain Monte Carlo (MCMC) simulations. The maximum clade credibility tree was calculated using TreeAnnotator from median heights, removing burn-in trees from the first 10% of the chains. The tree was plotted as previously described, with posterior probability rounded to the thousandth and age estimates plotted using ggplot(Wickham 2016) and ips(Heibl 2008).

### Pyseer analysis

To see if we could find any gene enrichment in *Tannerella forsythia M*AGs from Oceania, we ran pyseer v1.3.10(Lees et al. 2018) using the fixed model, correcting for population structure. We compared all MAGs from Oceania (n=25) to the MAGs from all other locations (n=85). P-value < 0.02 was considered significant, and effect size > 0.5 was considered large. We created a Manhattan plot to visualise the results using ggplot(Wickham 2016).

## Data Availability

All data used in this study comes from prior publications and is publicly available. All data accessions used in this study are listed in Supplementary Tables S1 and S2.

## Acknowledgments

We thank Maxime Borry for suggestions on GWAS approaches, Adam Ben Rohrlach for discussions on statistical approaches, and Alexander Hübner for assistance with MAG quality evaluation. This research was supported by the Werner Siemens Stiftung (“Paleobiotechnology” to C.W.), the Deutsche Forschungsgemeinschaft (DFG, German Research Foundation) under Germanýs Excellence Strategy (EXC 2051 – Project-ID 390713860 to C.W.), Carolyn Weinberg and the Radcliffe Institute for Advanced Study, the American School of Prehistoric Research, and the Max Planck-Harvard Research Center for the Archaeoscience of the Ancient Mediterranean (MHAAM).

## Supplementary tables and figures

## Tables

Supplementary Table S1. Metadata for Tannerella MAG and isolate genomes.

Supplementary Table S2. Metadata for Porphyromonas MAG and isolate genomes.

Supplementary Table S3. Virulence factors function and annotations.

Supplementary Table S5. Genomes used as training files for Prokka annotations.

Supplementary Table S4. fimA genotyping of MAGs.

Supplementary Table S6. Pyseer summary table.

## Figures

**Supplementary Figure S1.**
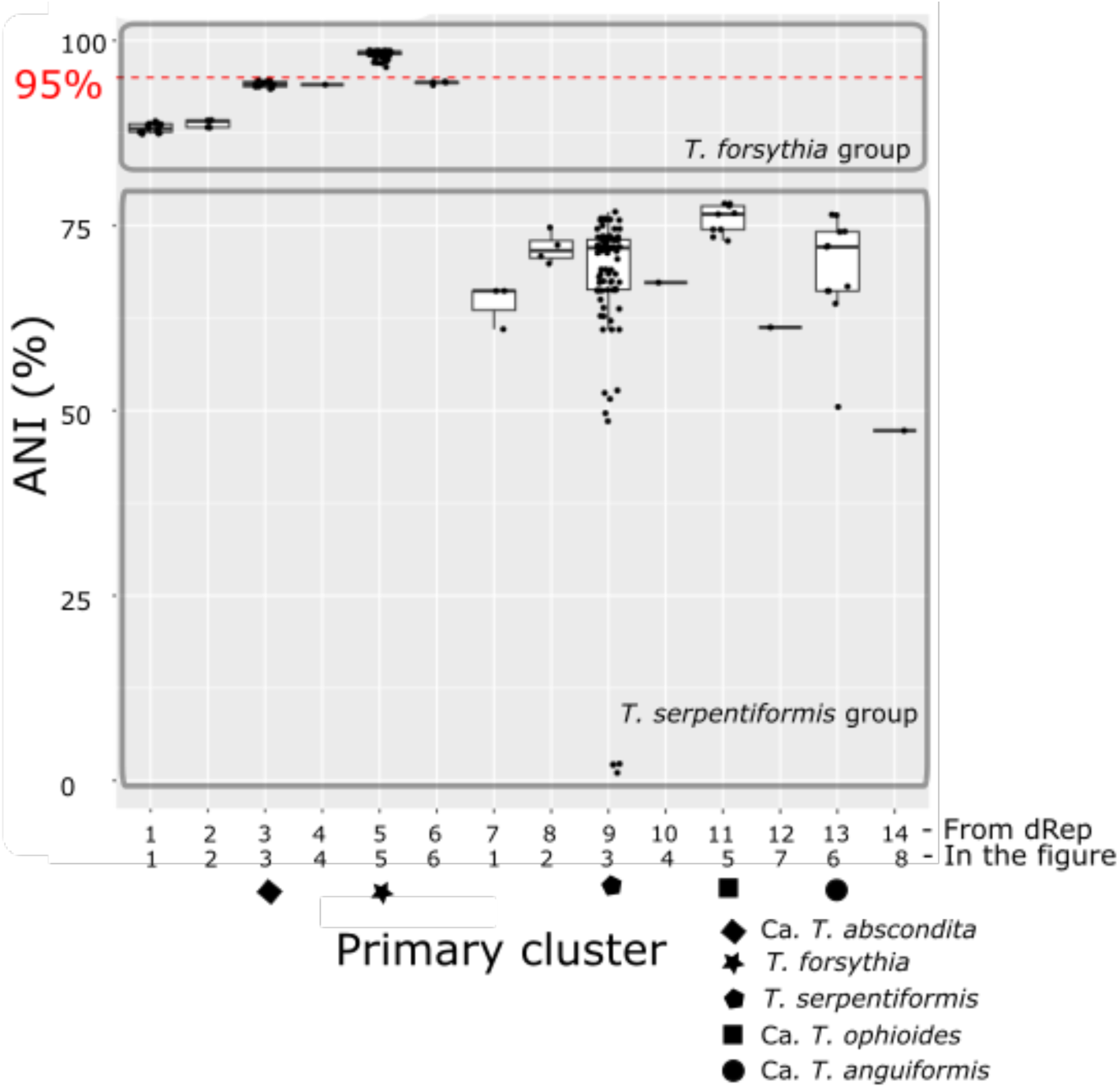
Average nucleotide identity (ANI) between all *Tannerella* MAGs and the *Tannerella forsythia* MAGs. MAGs with > 95% ANI (represented in the dashed red line) are considered to be a single species (in this figure *T. forsythia*), while MAGs falling below this cut-off are placed in independent clusters. The primary cluster numbers here correspond to those shown in the main text figures. See Supplementary table S1 to see the correspondence between the original cluster number generated by dRep and the updated main text cluster number. The three MAGs in the *T. serpentiformis* plot with ANI values below 25% are from howler monkeys.

**Supplementary Figure S2.**
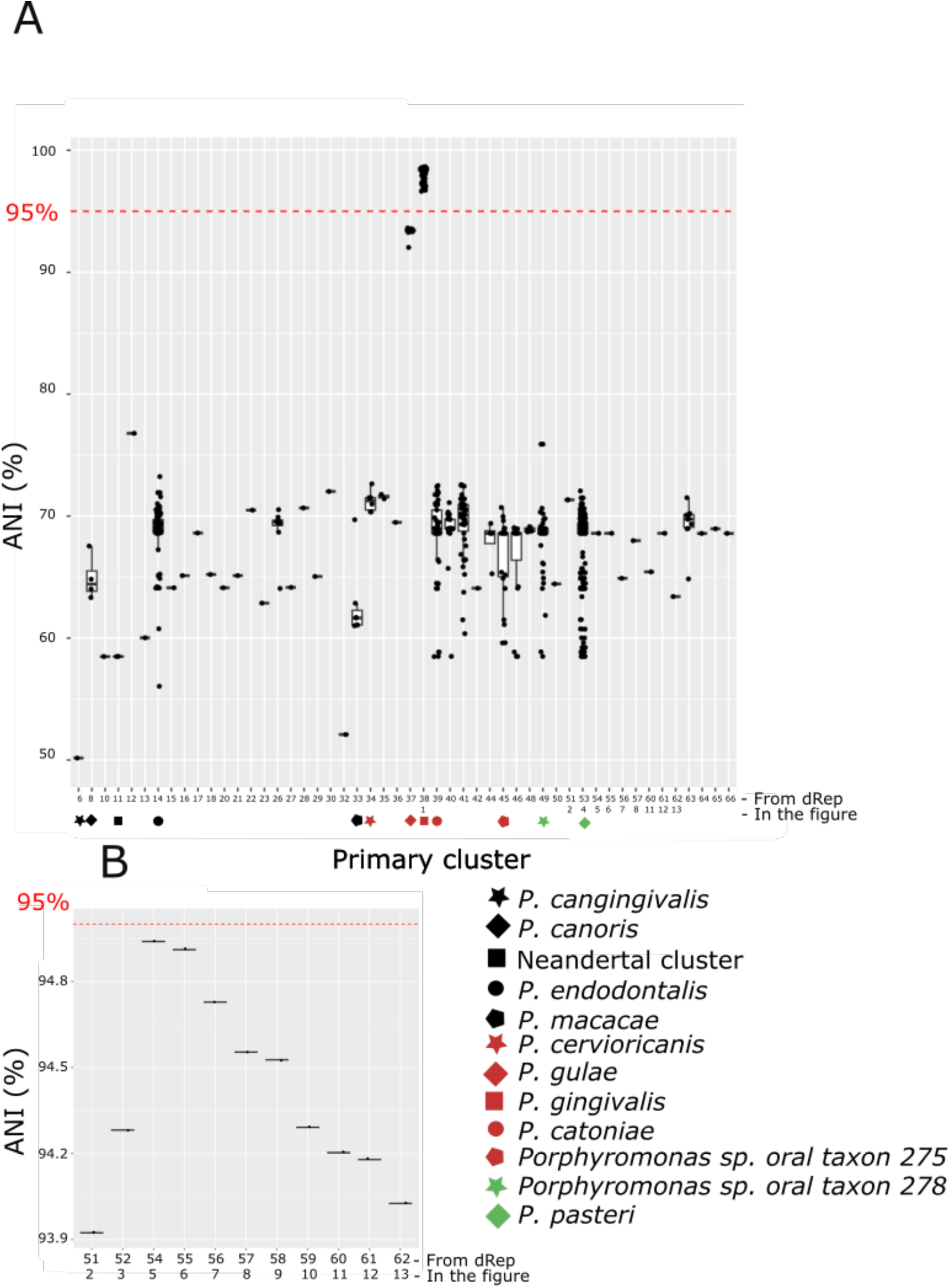
Average nucleotide identity (ANI) between all *Porphyromonas* MAGs. MAGs with > 95% ANI (represented in the dashed red line) are considered to be a single species (**A.** *P. gingivalis*; **B.** *P. pasteri*), while MAGs falling below this cut-off are placed in independent clusters. Only similarities > 50% are shown. The primary cluster numbers here correspond to those shown in the main text figures. See Supplementary Tables S1 and S2 to see the correspondence between the original cluster number generated by dRep and the updated main text cluster number. **A.** ANI of *Porphyromonas* MAGs in each cluster against the *P. gingivalis* cluster MAGs. The star indicates *P. pasteri*. **B.** All the *P. pasteri* cluster MAGs against the *P. pasteri* unique primary cluster. These MAGs falling outside the 95% ANI cut-off may have lower completeness/contamination that prevents them from reaching 95% ANI with other *P. paster*i MAGs.

**Supplementary Figure S3.**
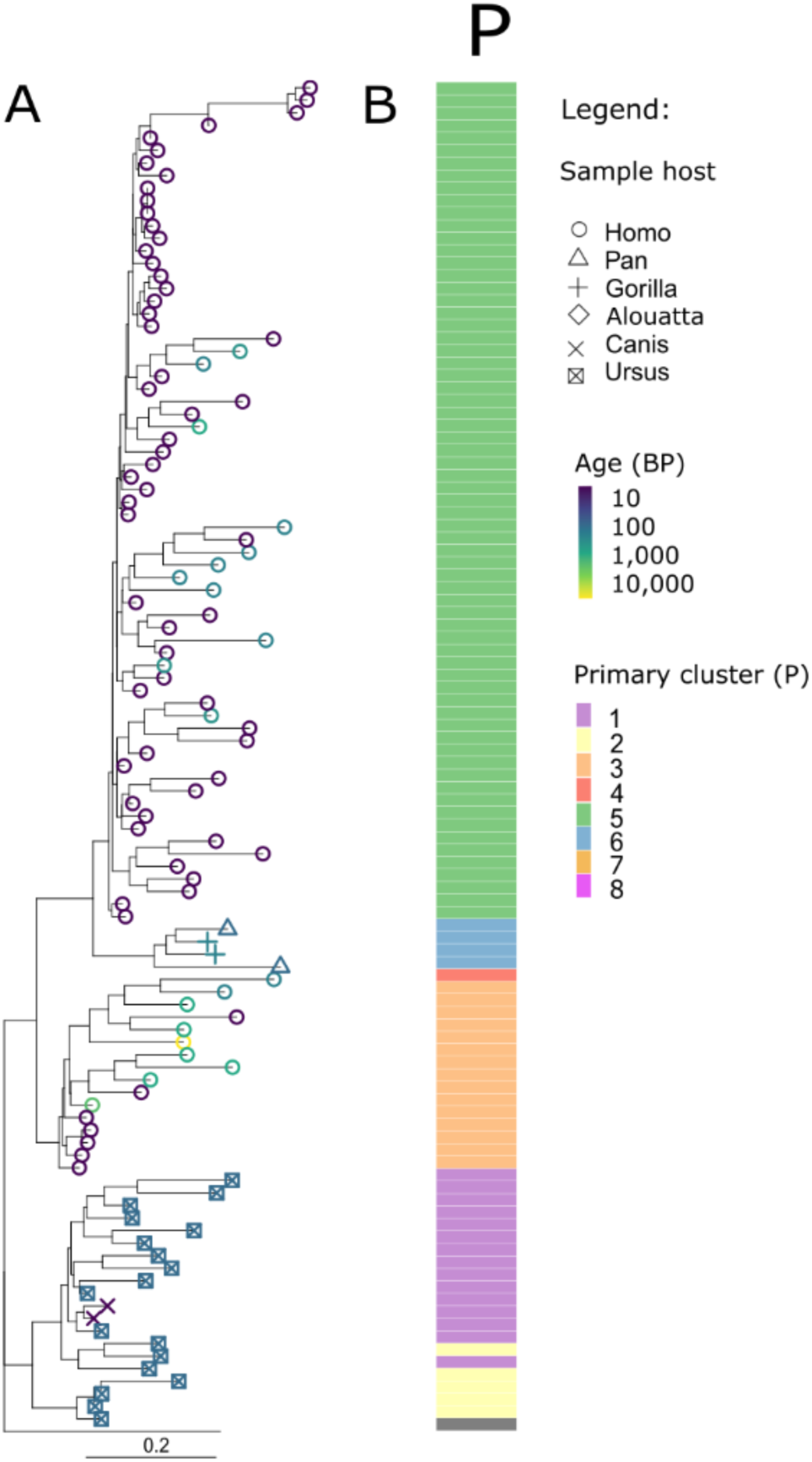
Phylogenetic relationship of *Tannerella forsythia* cluster MAGs using 400 universal genes from PhyloPhlan to confirm the tree structure. The tree tip color corresponds to the sample’s age in years before present (BP), and the tip shape to the sample host. The matrix gives information about the species cluster (based on 95% ANI cut-offs). The metadata matrix displays the species clusters with an average nucleotide identity (ANI) of ≥ 95%. The scale bar indicates substitution per position. *Tannerella serpentiformis* was used as the outgroup to root the tree. **A.** Maximum-likelihood tree. **B.** Primary cluster metadata matrix.

**Supplementary Figure S4.**
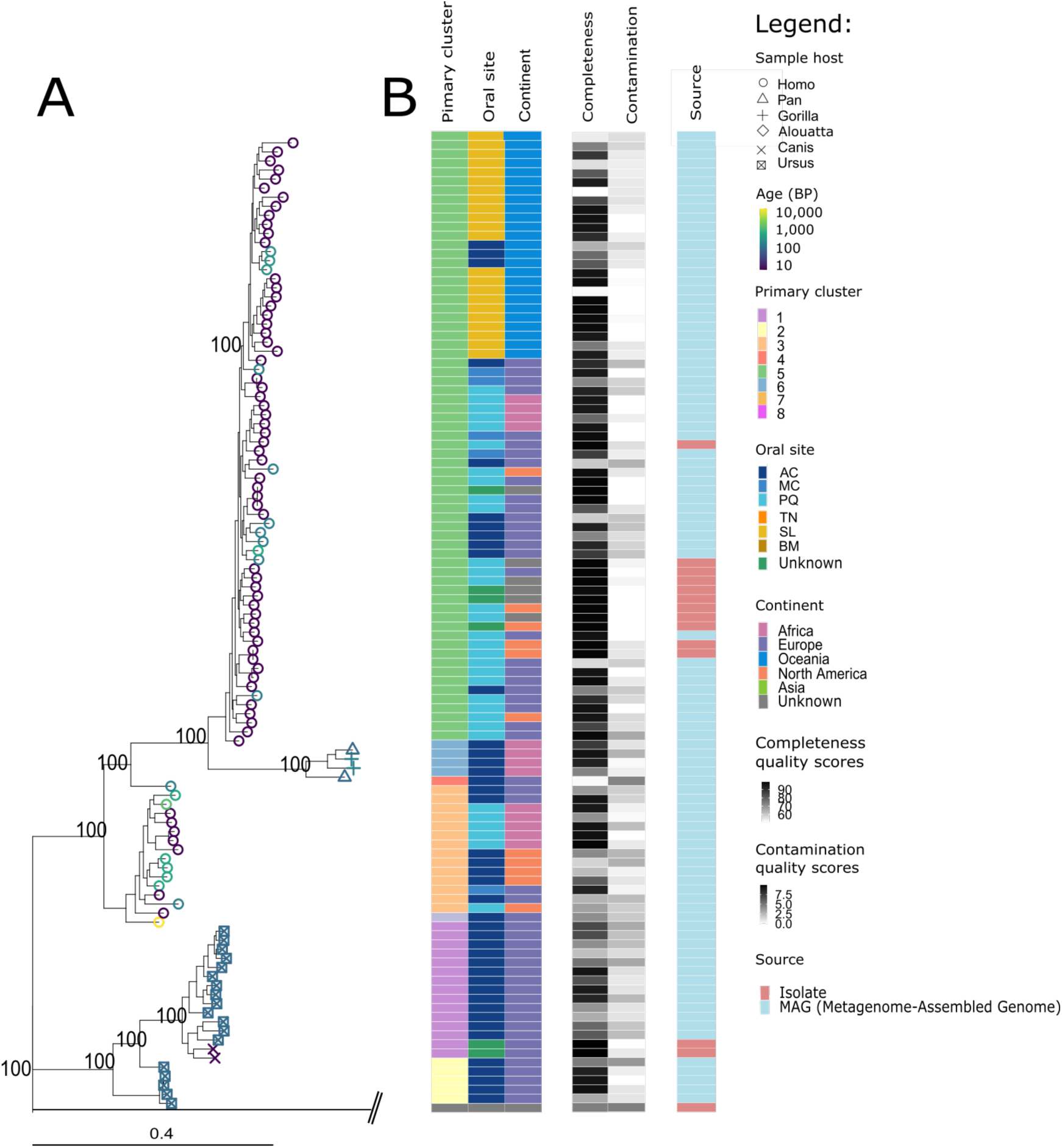
Phylogenetic relationship of *Tannerella forsythia* MAGs to the sample and MAG metadata. The tree tip color corresponds to the sample’s age in years before present (BP), and the tip shape to the sample host. The matrix gives information about the species cluster (based on 95% ANI cut-offs), oral site, continent, and original publication of the samples from which MAGs are assembled, and completeness and contamination estimates of the MAGs. Bootstrap support based on 100 replicates is indicated as percentages for principal branches. The scale bar indicates substitution per position. *Tannerella serpentiformis* was used as the outgroup to root the tree. **A.** Maximum-likelihood phylogenetic tree of *Tannerella forsythia* group (same as main Figure 2A). **B**. Metadata matrix of *Tannerella forsythia* group. Oral site - Oral: The sampling site was not specified. AC: Ancient calculus, MC: modern calculus, PQ: plaque, TN: tongue, SL: saliva, BM: buccal mucosa, HP: hard palate.

**Supplementary Figure S5.**
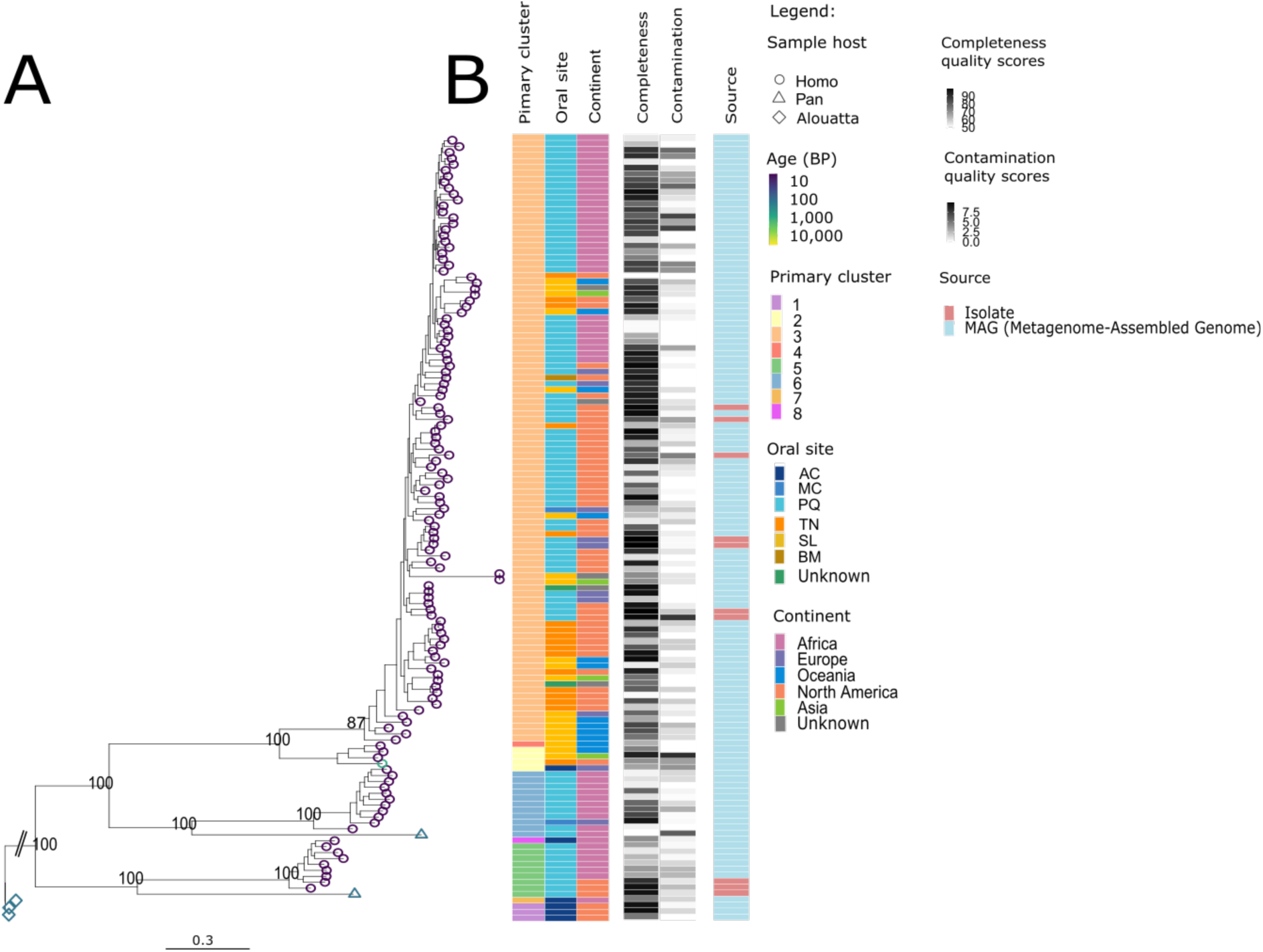
Phylogenetic relationship of *Tannerella serpentiformis* to geographic origin, oral sample site, studies, completeness, and contamination estimates of MAGs. The tree tip color corresponds to the sample’s age in years before present (BP), and the tip shape to the sample host. The metadata matrix gives information about the species cluster (based on 95% ANI cut-offs), oral site, continent, and original publication of the samples from which MAGs are assembled, as well as completeness and contamination estimates of the MAGs. Bootstrap support values based on 100 replicates are indicated as percentages for principal branches. The scale bar indicates substitutions per position. The tree was rooted using a MAG from a howler monkey (*Alouatta*) sample. **A.** Maximum-likelihood phylogenetic tree of *Tannerella serpentiformis* group. **B.** Metadata matrix of *Tannerella serpentiformis* group. Oral site - Oral: The sampling site was not specified. AC: Ancient calculus, MC: modern calculus, PQ: plaque, TN: tongue, SL: saliva, BM: buccal mucosa, HP: hard palate.

**Supplementary Figure S6.**
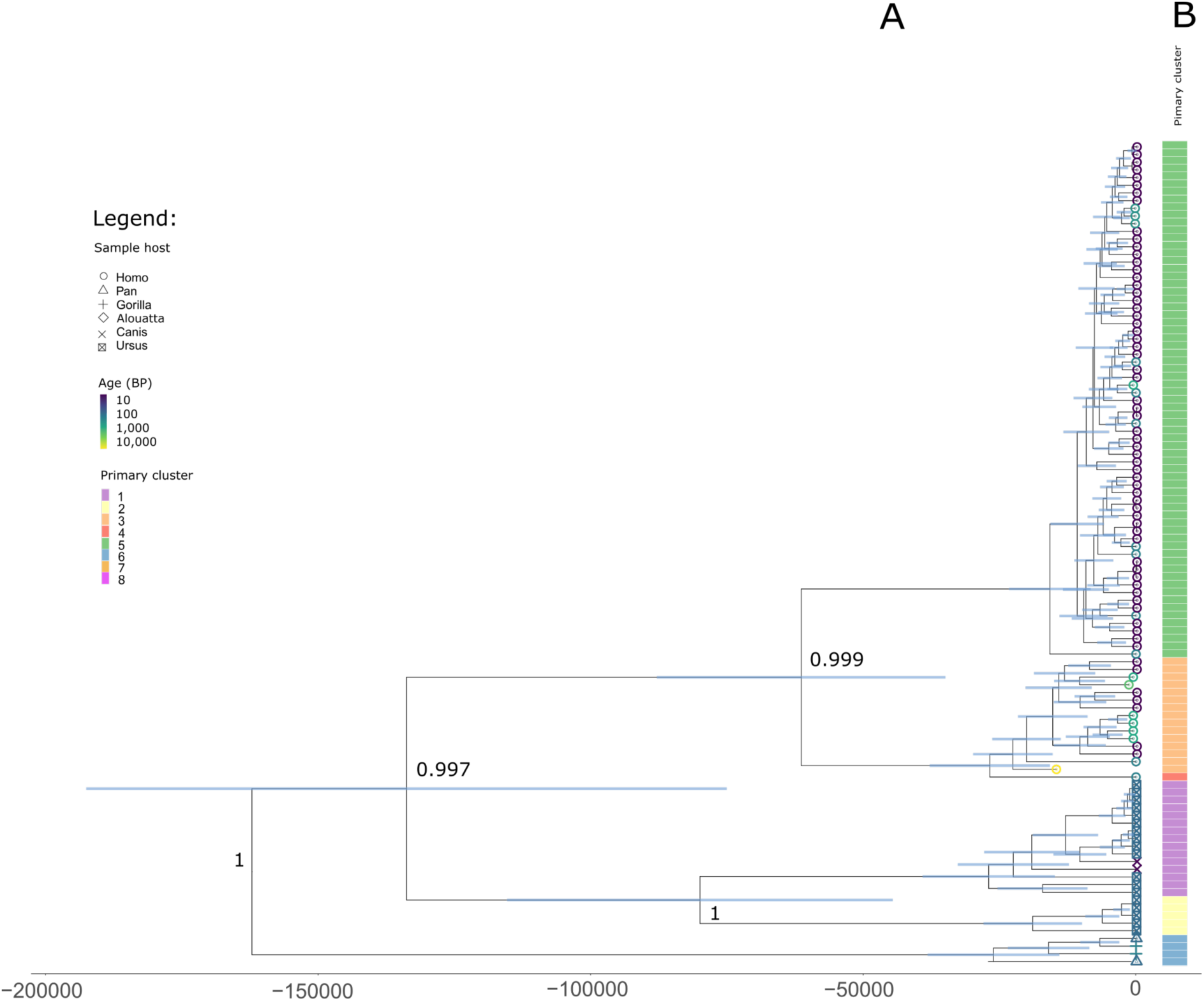
BEAST analysis to estimate the split time between *T. forsythia* and Ca. *T. abscondita*. Tree tip color corresponds to sample age in years before present (BP), and the tip shape to the sample host. The metadata matrix shows the 95% ANI-based species cluster. Posterior probabilities are indicated for principal branches. The blue bars indicate the 95% confidence interval of the age estimates. **A**. Maximum likelihood BEAST tree of *Tannerella forsythia* group. **B**. Metadata matrix of *Tannerella forsythia* group.

**Supplementary Figure S7.**
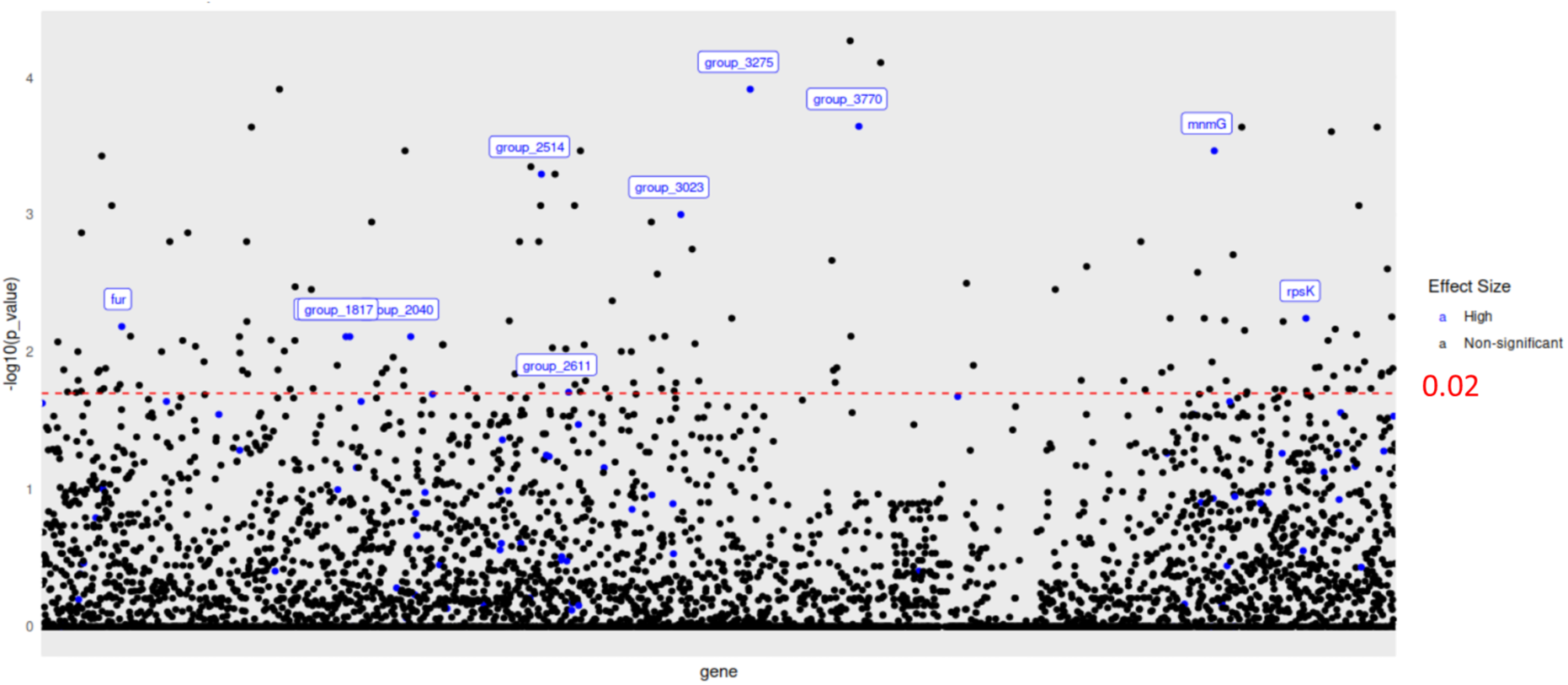
Gene association with sample geographic origin. Gene Manhattan plot from pyseer. Analysis used a fixed model, with p-values corrected for population structure and for false positives using the Benjamini-Hochberg correction. Dots are labeled and colored blue if the p-value is less than 0.02 and the effect size is high (>0.5), indicating a significant association with the continent of origin of the sample. All the MAGs are from the *Tannerella forsythia* clade; 25 Oceanian MAGs are compared to 85 MAGs/reference genomes from other parts of the world.

**Supplementary Figure S8.**
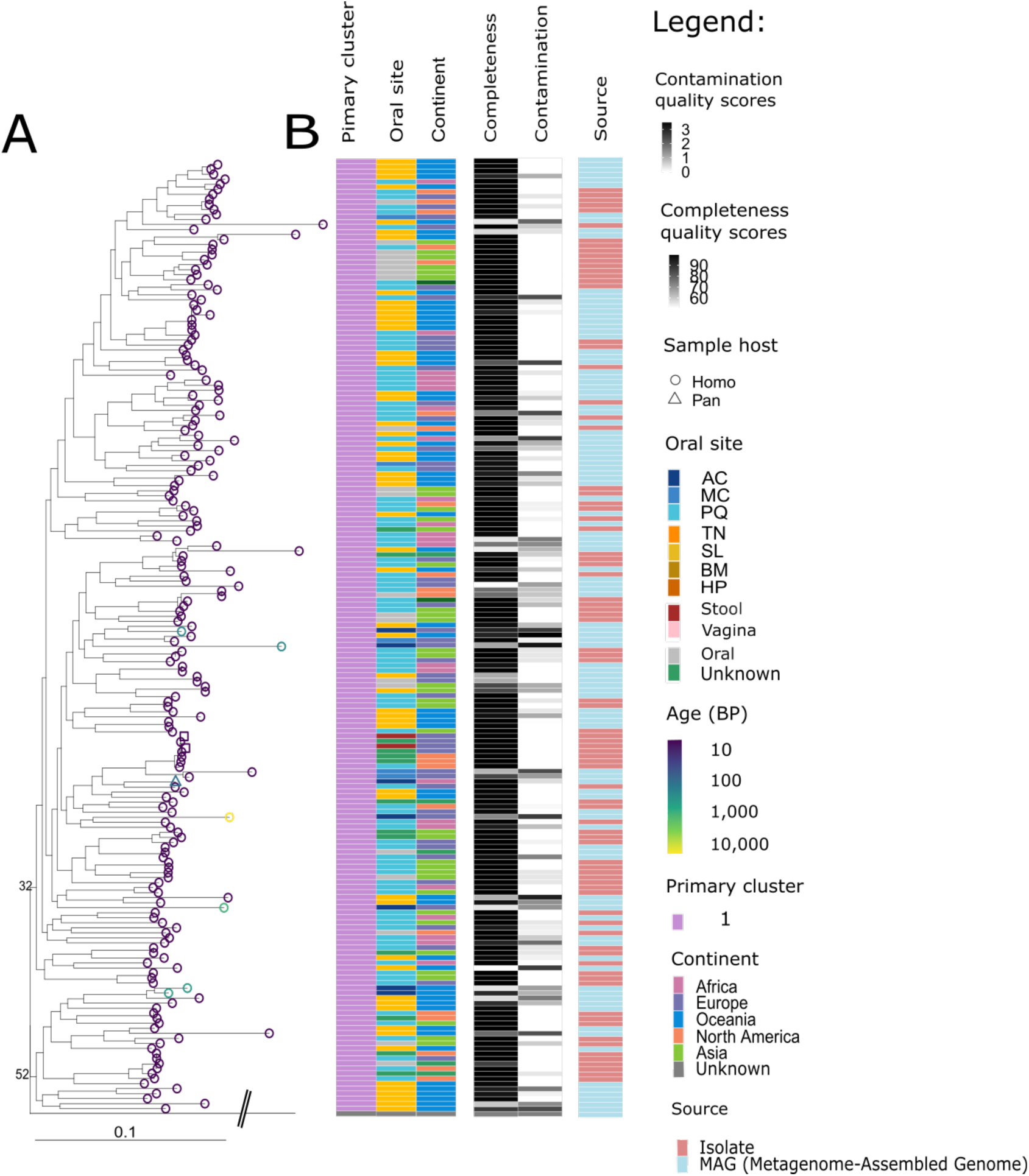
Phylogenetic relationship of *Porphyromonas gingivalis* to geographic origin, oral sample site, studies, completeness, and contamination estimates of MAGs. The tree tip color corresponds to the sample’s age in years before present (BP), and the tip shape to the sample host. The matrix gives information about the species cluster (based on 95% ANI cut-offs), oral site, continent, and original publication of the samples from which MAGs are assembled, and completeness and contamination estimates of the MAGs. The tree was rooted using a *Porphyromonas pasteri* reference genome from NCBI. **A**. Maximum-likelihood phylogenetic tree of *Porphyromonas gingivalis* species cluster. **B**. Metadata matrix of *Porphyromonas gingivalis*. Oral site - Oral: The sampling site was not specified. AC: Ancient calculus, MC: modern calculus, PQ: plaque, TN: tongue, SL: saliva, BM: buccal mucosa, HP: hard palate.

**Supplementary Figure S9.**
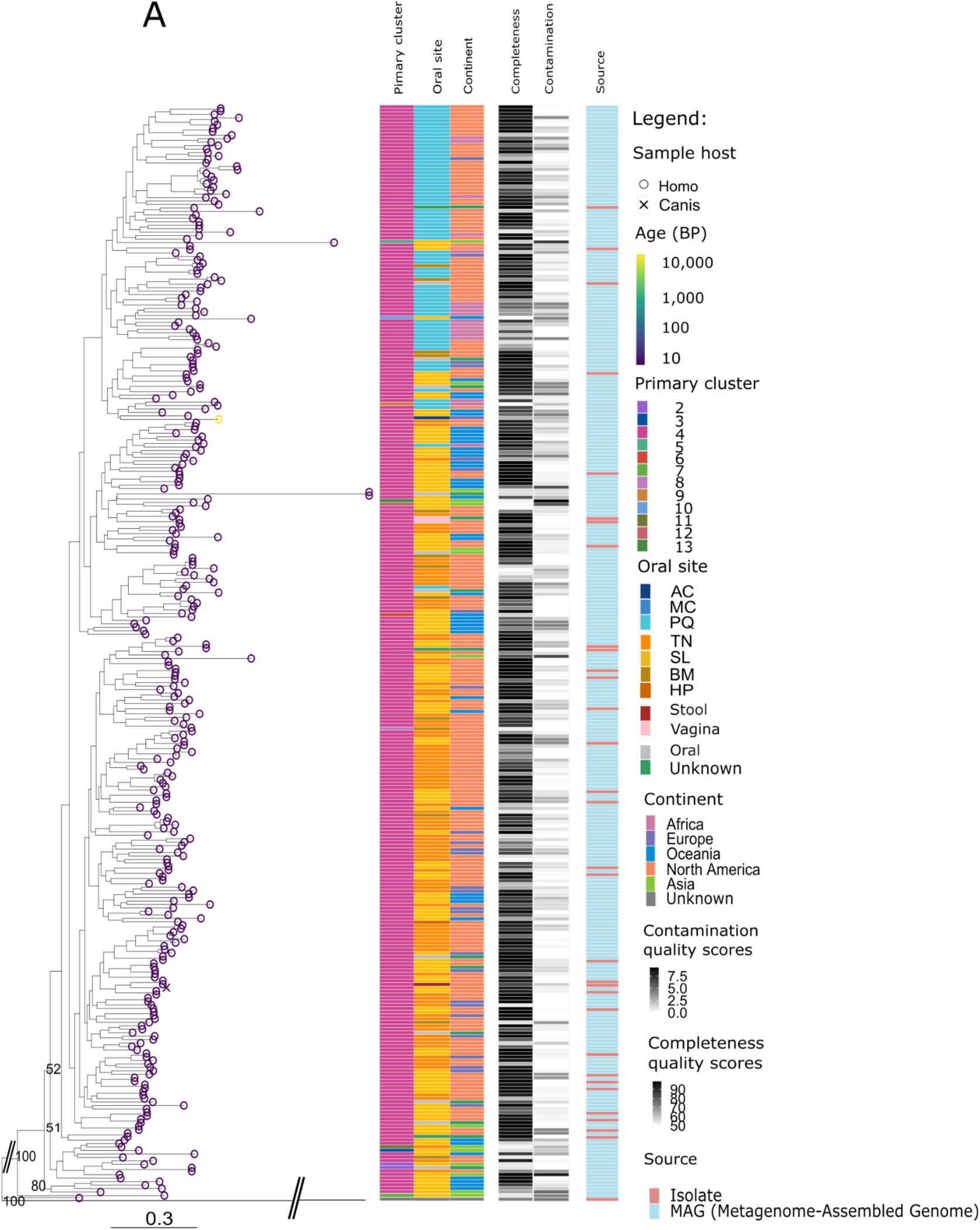
Phylogenetic relationship of *Porphyromonas pasteri* to geographic origin, oral sample site, studies, completeness, and contamination estimates of MAGs. The tree tip color corresponds to the sample’s age in BP (years Before Present), and the tip shape to the sample host. The matrix gives information about the species cluster (based on 95% ANI cut-offs), oral site, continent, and original publication of the samples from which MAGs are assembled, and completeness and contamination estimates of the MAGs. The tree was rooted using the midpoint option in R. A. Maximum-likelihood phylogenetic tree of *Porphyromonas pasteri* species cluster. B. Metadata matrix. Oral site - Oral: The sampling site was not specified. AC: Ancient calculus, MC: modern calculus, PQ: plaque, TN: tongue, SL: saliva, BM: buccal mucosa, HP: hard palate.

**Supplementary Figure S10.**
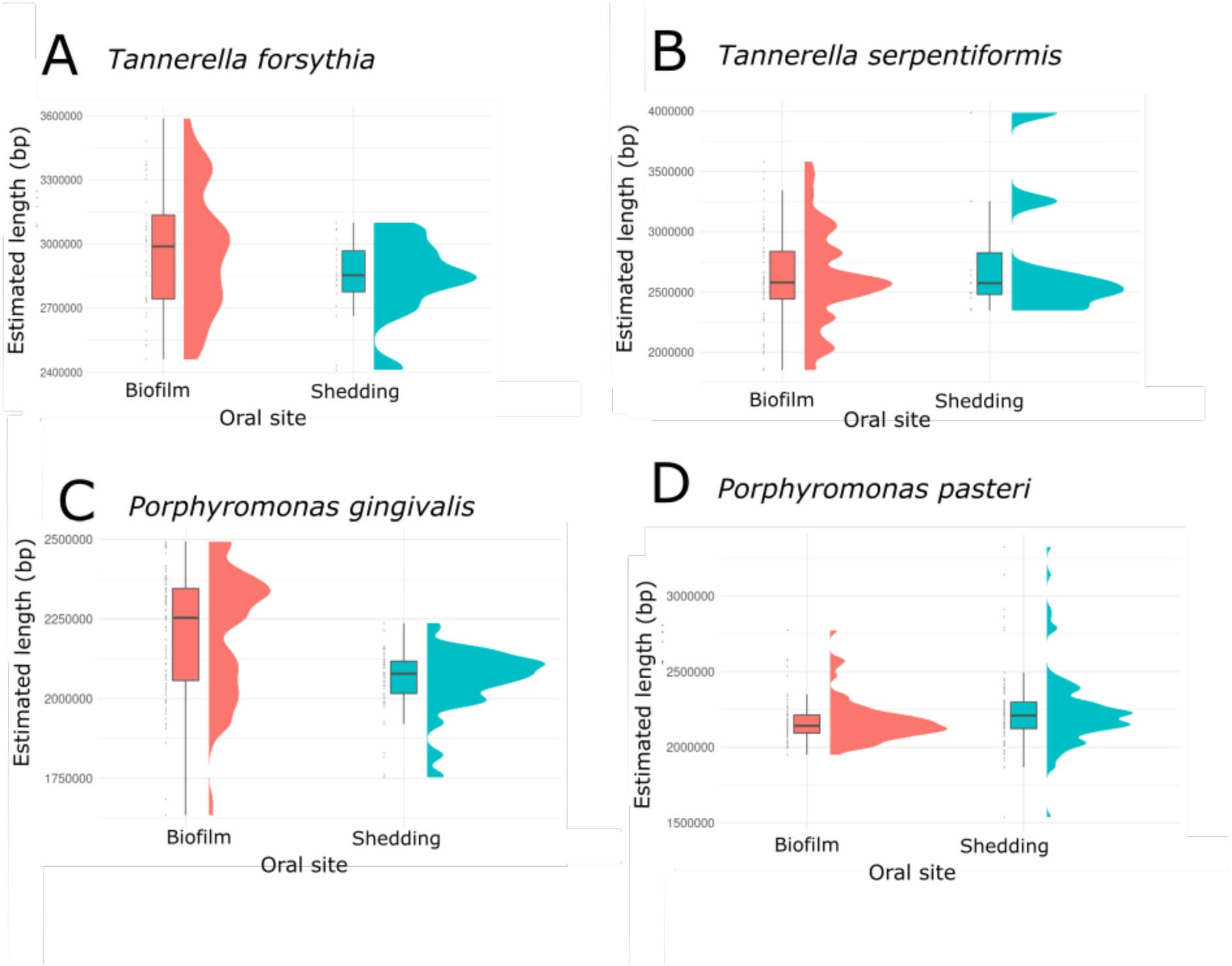
Genome length comparison between shedding and non-shedding biofilm surfaces. The length is estimated based on the completeness of the MAGs and the length of the corresponding reference genome from NCBI RefSeq. **A.** *T. forsythia*. **B**. *T. serpentiformis*. **C**. *P. gingivalis*. **D**. *P. pasteri.* No significant differences were detected between MAGs from biofilms vs. shedding surfaces. Shedding surfaces include saliva, tongue, buccal mucosa, and hard palette; non-shedding biofilm surfaces include dental plaque and calculus (ancient and modern).

